# State-dependent modulation of spiny projection neurons controls levodopa-induced dyskinesia in a mouse model of Parkinson’s disease

**DOI:** 10.1101/2025.01.02.631090

**Authors:** Shenyu Zhai, Qiaoling Cui, David Wokosin, Linqing Sun, Tatiana Tkatch, Jill R. Crittenden, Ann M. Graybiel, D. James Surmeier

## Abstract

In the later stages of Parkinson’s disease (PD), patients often manifest levodopa-induced dyskinesia (LID), compromising their quality of life. The pathophysiology underlying LID is poorly understood, and treatment options are limited. To move toward filling this treatment gap, the intrinsic and synaptic changes in striatal spiny projection neurons (SPNs) triggered by the sustained elevation of dopamine (DA) during dyskinesia were characterized using electrophysiological, pharmacological, molecular and behavioral approaches. Our studies revealed that the intrinsic excitability and functional corticostriatal connectivity of SPNs in dyskinetic mice oscillate between the on- and off-states of LID in a cell- and state-specific manner. Although triggered by levodopa, these rapid oscillations in SPN properties depended on both dopaminergic and cholinergic signaling. In a mouse PD model, disrupting M1 muscarinic receptor signaling specifically in iSPNs or deleting its downstream signaling partner CalDAG-GEFI blunted the levodopa-induced oscillation in functional connectivity, enhanced the beneficial effects of levodopa and attenuated LID severity.

## Introduction

In Parkinson’s disease (PD), the loss of the dopaminergic neurons in the substantia nigra pars compacta (SNc) disrupts basal ganglia circuitry, leading to bradykinesia, rigidity and tremor ^1^. In the early stages of PD, systemic levodopa administration boosts the production and release of dopamine (DA), effectively ameliorating motor symptoms. However, as the disease progresses, the levodopa dose needed to achieve symptomatic benefits rises and the machinery regulating extracellular DA wanes in efficacy ^2–5^. As a result, DA signaling is dysregulated, rising dramatically for hours after levodopa treatment and then falling to very low levels until the next dose is taken.

This abnormal, slow oscillation in brain DA concentration triggers alterations in basal ganglia circuits that result in uncontrolled movements (i.e., dyskinesia) shortly after levodopa is taken. Although several parts of the basal ganglia have been implicated in the emergence of levodopa-induced dyskinesia (LID), there is a consensus that the striatum is a critical site of pathophysiology6,7.

The principal neurons of the striatum are GABAergic spiny projection neurons (SPNs), which constitute ∼90% of all striatal neurons. About half of SPNs – so-called direct pathway SPNs (dSPNs) – project directly to the output nuclei of the basal ganglia, promoting action selection. The other half – the indirect pathway SPNs (iSPNs) – project to the external segment of the globus pallidus and thus are indirectly connected to the output nuclei; activity in iSPNs is generally thought to suppress contextually inappropriate actions ^8,9^. Due to their differential expression of DA receptors, iSPNs and dSPNs are modulated by DA in opposite ways. In dSPNs, G_s/olf_-coupled D1 DA receptors (D1Rs) stimulate adenylyl cyclase (AC) and protein kinase A (PKA), increasing intrinsic excitability, enhancing glutamatergic synaptic transmission, and facilitating long-term synaptic potentiation (LTP). In contrast, in iSPNs, G_i_-coupled D2 DA receptors (D2Rs) inhibit AC and stimulate phospholipase C, decreasing intrinsic excitability, attenuating glutamatergic transmission, and promoting long-term synaptic depression (LTD) ^9–13^. Importantly, in the healthy striatum, dopaminergic signaling is episodic, being linked to the initiation of actions and to the outcomes of actions ^14^. These transient signaling events are thought to be critical to connecting actions and their outcomes to modifications in axospinous synaptic strength that underlie the acquisition of habits and contextually appropriate goal-directed actions ^10,14^.

The impact of dopaminergic signaling on SPNs is normally modulated by a dynamic interaction with autonomously active, giant, cholinergic interneurons (ChIs). Acting through D2Rs, striatal DA release inhibits the autonomous spiking of ChIs and their release of acetylcholine (ACh) ^15–18^. The ACh released by ChIs acts on iSPNs and dSPNs in ways that counter those of DA. In dSPNs, which primarily express G_i_-coupled M4 muscarinic receptors (M4Rs), ACh signaling blunts the effects of D1R activation and promotes LTD induction ^19–21^, whereas, in iSPNs, which only express M1 muscarinic receptors (M1Rs), ACh signaling enhances somatic excitability, dendritic integration and LTP induction ^22–24^. The M1R-mediated modulation of iSPN dendrites is dependent upon CalDAG-GEFI (CDGI), a striatum-enriched, Ca^2+^-activated guanine nucleotide exchange factor ^22,25^. Thus, the interaction between dopaminergic and cholinergic signaling not only modulates the moment-to-moment excitability of striatal ensembles coordinating purposeful movement, but also the long-term changes in synaptic strength that guide future behavior.

In models of late-stage PD, where most of dopaminergic neurons innervating the striatum have been lost, there appears to be a sustained enhancement of ACh release by ChIs ^26^. This shift is attributable to an elevation in ChI intrinsic excitability, a strengthening of their excitatory glutamatergic input from the parafascicular nucleus (PFN), and a dis-inhibition of ACh release from ChI terminals ^27–30^. In animal models of PD, optogenetic or chemogenetic inhibition of ChIs alleviates motor deficits ^21,29,31^. Furthermore, optogenetic activation of ChIs in healthy mice induces a parkinsonian-like state ^32^. The hypothesis that elevated ACh release contributes to the hypokinetic features of PD is also supported by the clinical observation that muscarinic receptor antagonists are of symptomatic benefit ^33^. But, the involvement of ChIs in the dyskinesia induced by levodopa treatment is controversial ^34–36^. On one hand, boosting cholinergic signaling appears to attenuate LID. For example, enhancing M4R signaling in dSPNs attenuates LID severity ^20,37^. On the other hand, several studies suggest that ChI ablation or disruption of cholinergic signaling attenuates LID severity ^38,39^. One critical gap in these studies is an assessment of how cholinergic signaling and SPNs are changing between the period when mice are dyskinetic and striatal DA is high (on-state) and when mice are hypokinetic and striatal DA is low (off-state). It is highly likely that the activity of ChIs and cholinergic signaling in these two states are very different. Moreover, as DA and ACh normally work in concert to control long-term changes in the functional connectomes of SPN underlying learning, it could be that the aberrant interaction between these neuromodulators induced by high doses of levodopa lead to pathological ‘learning’ and alterations in circuitry that carry over from one state to the next.

To help fill this fundamental gap in our understanding, a combination of electrophysiological, imaging, genetic, pharmacological and behavioral approaches were employed in a mouse model of LID. These studies revealed that LID on- and off-states were associated with bidirectional, cell-type specific changes in intrinsic excitability and synaptic connectivity. In addition, these studies demonstrated that ACh release by ChIs was elevated in the parkinsonian and LID off-states. Moreover, the release of ACh by ChIs continued to be negatively modulated by D2Rs in tissue from dyskinetic mice, arguing that striatal cholinergic signaling was inhibited in the on-state. The dysregulation of cholinergic signaling was critical to the state-dependent alterations in synaptic strength and dendritic spine architecture in iSPNs. Blunting the impact of ChIs on iSPNs not only attenuated dyskinetic behaviors, it enhanced the beneficial effects of levodopa treatment.

## Results

### Intrinsic excitability and synaptic connectivity of dSPNs increased in the on-state of LID

Previous studies of how the induction of LID in rodents alters the intrinsic excitability and synaptic connectivity of SPNs have focused on the ‘off-state’ (usually 24-48 hours after the last administration of levodopa) ^40–43^. The implicit assumption of these studies is that while SPN properties change over the days during the induction of LID, they do not change in the relatively short period after termination of levodopa treatment. To test this hypothesis, the properties of dSPNs were compared in the off-state and on-state of LID. *Drd1*-Tdtomato bacterial artificial chromosome (BAC) transgenic mice were rendered parkinsonian by unilateral medial forebrain bundle (MFB) injection of 6-OHDA (Fig. 1A). Three to four weeks later, the extent of DA denervation was assessed using a drug-free, forelimb-use asymmetry test (also called cylinder test). Mice with a near-complete lesion were given dyskinesiogenic doses of levodopa every other day for at least five sessions (6 mg/kg for the first two sessions and 12 mg/kg for later sessions, supplemented with 12 mg/kg benserazide) ^4,20,40^. Mice were then sacrificed either 30 min after the last levodopa dose (on-state) or 24-48 hours after the last dose (off-state). In the on-state, phosphorylated ERK (pERK) in the DA-depleted striatum was elevated (Fig. 1B), as previously reported ^44,45^.

**Figure 1.**
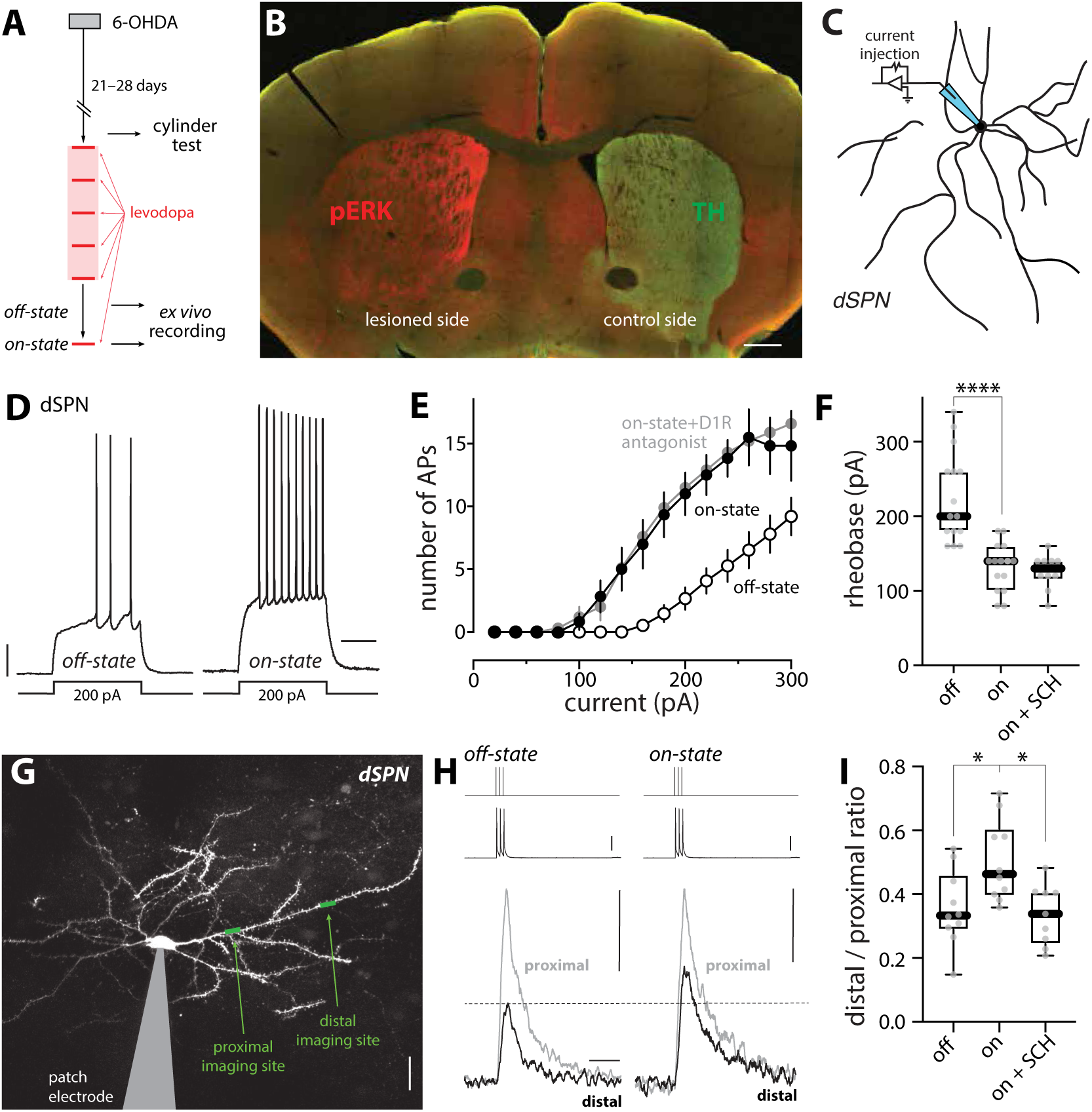
dSPN somatic and dendritic excitability changed between off- and on-states of LID. (A) Experimental timeline. Note that the LID off-state mice were sacrificed 24-48 hours following last levodopa injection for *ex vivo* experiments while LID on-state mice were sacrificed 30 min after the last levodopa injection. (B) Confocal image of a coronal brain slice from a unilateral 6-OHDA lesioned mouse immunostained with anti-phospho-ERK (pERK) antibody (red) and anti-tyrosine hydroxylase (TH) antibody (green). Scale bar is 0.5 mm. (C) Schematic illustrating the somatic excitability assay in dSPNs. (D) Sample voltage recordings from dSPNs in the off-state and on-state of LID in response to a 200- pA current injection (500 ms duration). Scale bars are 10 mV and 200 ms. (E) Current-response curves. Somatic excitability of dSPNs, as revealed by the number of action potentials (APs), was increased in the on-state and this increase was resistant to acute bath application of D1R antagonist SCH 23390 (3 μM). (F) Box plot summary of rheobase in dSPNs (off-state, n = 15 cells from 8 mice; on-state, n = 15 cells from 10 mice; on-state + SCH, n = 10 cells from 5 mice). **** p < 0.0001, Mann-Whitney test. (G) 2PLSM image of a whole-cell patched dSPN with imaging sites on proximal and distal dendrites indicated. Scale bar is 20 μm. (H) Current injections of 2 nA (top) are temporally aligned with corresponding somatic voltage changes (middle, scale bar is 25 mV) and fluorescence transients recorded from proximal and distal dendrites (bottom, scale bars are 0.1 ΔG/R_0_ and 0.5 s). (I) Box plot summary of dendritic excitability index (the distal to proximal ratio of area under the curve (AUC) of ΔG/R_0_) in dSPNs from off-state mice, on-state mice or on-state mice with acute bath application of SCH 23390 (off-state, n = 10 cells from 7 mice; on-state, n = 10 cells from 5 mice; on-state + SCH, n = 9 cells from 5 mice). * p < 0.05, Mann-Whitney test.

To assess somatic excitability, visually identified dSPNs in brain slices were subjected to whole-cell patch clamp recording ^20,40,46^ (Fig. 1C). The response to current steps was monitored and the relationship between step amplitude and evoked spiking plotted (Fig. 1D-E). In off-state dSPNs, the relationship between spike frequency and current intensity (i.e. F-I relationship) was similar to that described in previous studies ^40^. However, in the on-state, the somatic excitability of dSPNs was elevated: in on-state dSPNs, the rheobase current was significantly lower than that in the off-state (p < 0.0001) (Fig. 1F) and the number of spikes evoked by suprathreshold current steps was greater across a range of intensities (Fig. 1E). The shift in on-state somatic excitability was consistent with an elevation in on-state extracellular DA and activation of D1Rs on dSPNs ^47–52^. That said, in agreement with recent work suggesting that the D1R-mediated modulation of dSPN somatic excitability is persistent ^53^, bath application of the D1R antagonist SCH 23390 (3 µM) did not reverse the leftward shift in the F-I relationship curve or the reduction in rheobase (p = 0.589) (Fig. 1E-F).

Most of the surface area of SPNs is composed of active dendrites, making them key sites of dopaminergic modulation. To assess dSPN dendritic excitability in on- and off-states, a combination of patch clamp electrophysiology and two-photon laser scanning microscopy (2PLSM) was used. Specifically, dSPNs in *ex vivo* brain slices from off- or on-state mice were patch clamped, filled with Ca^2+^-sensitive dye Fluo-4 and Ca^2+^-insensitive dye Alexa Fluor 568 (to visualize dendrites), and injected with brief current steps (three 2 nA injections, 2 ms each, at 50 Hz). These somatically delivered steps evoked spikes that back-propagated into SPN dendrites ^23,54^. To assess the spread of back-propagating action potentials (bAPs), 2PLSM was used to determine evoked changes in Fluo-4 fluorescence along dendrites produced by transient opening of voltage-dependent Ca^2+^ channels ^23,54^. The magnitudes of the Ca^2+^ signals at proximal (∼40 µm from soma) and distal (∼90 µm from soma) dendritic locations served as a surrogate estimate of the extent of dendritic depolarization produced by the bAPs (Fig. 1G, H). To generate an estimate of bAP invasion that was independent of dye concentration, laser intensity and other experimental variables, the bAP-evoked Ca^2+^ signal in a distal dendritic segment was normalized by the bAP-evoked Ca^2+^ signal in the proximal region of the same dendrite. This index of dendritic excitability was significantly greater in on-state dSPNs than in off-state dSPNs (p = 0.0115) (Fig. 1I), as expected from an elevation in D1R signaling ^51,52^. However, unlike somatic excitability, the elevation in on-state dSPN dendritic excitability was readily reversed by bath application of the D1R antagonist SCH 23390 (p = 0.0133) (Fig. 1I).

In addition to modulating intrinsic excitability, dopaminergic signaling regulates the strength of glutamatergic synapses on dSPNs ^10,55^. Activation of D1Rs promotes the induction of LTP at axospinous, corticostriatal synapses, which can be manifested as a structural enlargement of spines ^56,57^. To determine LID-associated structural changes in dendritic spines, dSPNs in *ex vivo* brain slices were loaded with Alexa Fluor 568 through the patch pipette and then their dendritic morphology optically dissected using 2PLSM (Fig. 2A) ^40^. Consistent with previous reports ^40–42,58^, dSPN spine density was not altered in either proximal or distal dendrites 3-4 weeks following unilateral 6-OHDA lesion (p = 0.654 and 0.756 for distal and proximal dendrites, respectively) (Fig. 2, A-B). However, in dSPNs taken from off-state LID mice, spine density in both dendritic regions was significantly reduced (p < 0.0001 in both proximal and distal dendrites) (Fig.2, A-B). In dSPNs taken from on-state LID mice, spine density was not significantly different from that in the off-state (distal, p = 0.115; proximal, p = 0.106) (Fig. 2, A-B). However, in the on-state, enlarged, mushroom spines were more common in both distal and proximal dendrites (on-state vs. off-state: distal, p = 0.0025; proximal, p = 0.0184) (Fig. 2C).

**Figure 2.**
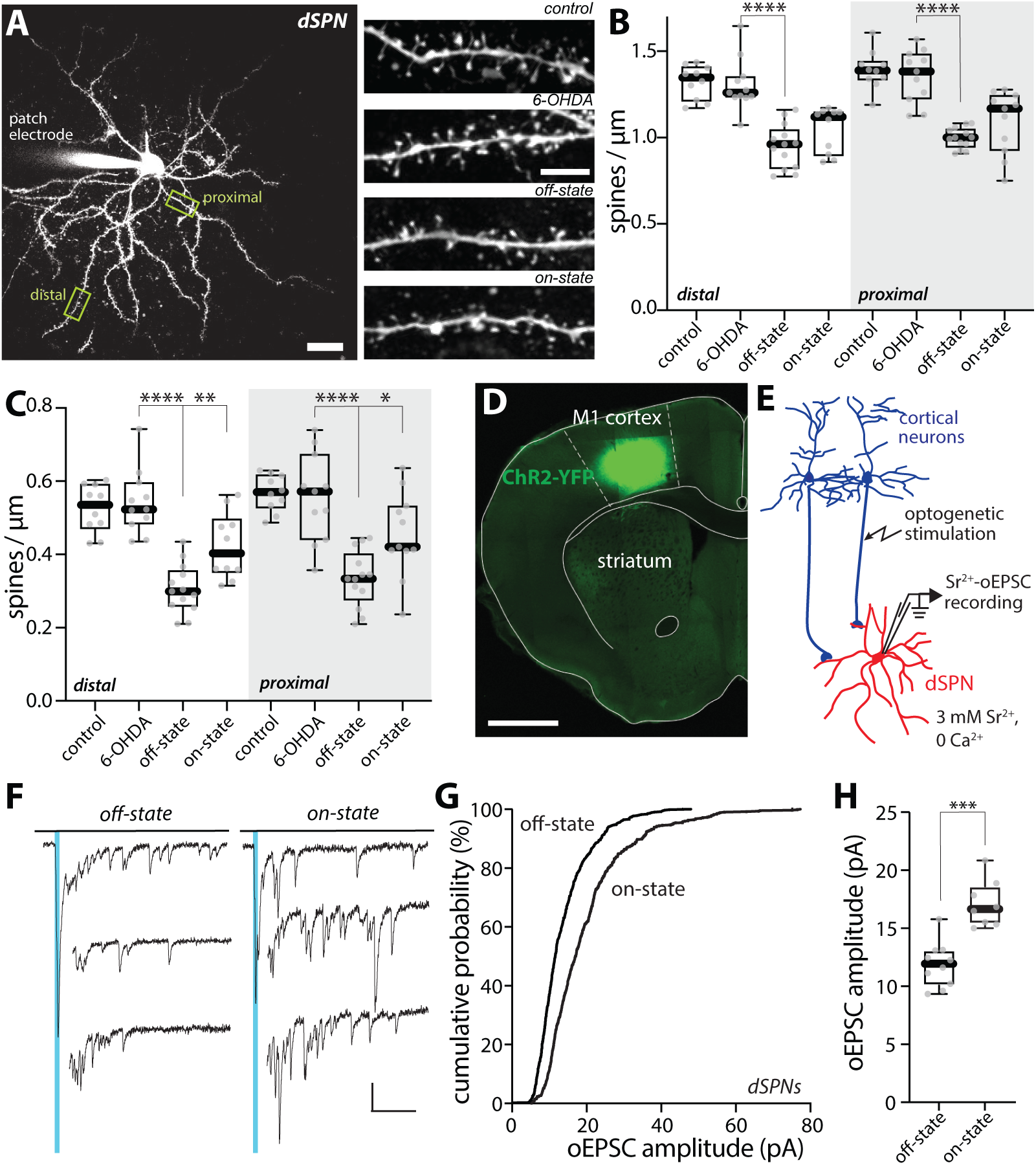
dSPN spine morphology and unitary synaptic strength changed between off- and on-states of LID. (A) Left, a low-magnification 2PLSM image of a patched dSPN with its dendritic tree visualized by Alexa dye and its proximal and distal locations used for spine density measurements delineated. Scale bar is 20 μm. Right, sample 2PLSM images of dendritic segments of dSPNs from control, 6-OHDA lesioned, LID off-state and LID on-state mice. Scale bar represents 5 μm. (B-C) Box plot summaries of density of all dendritic spines (B) and density of mushroom-type spines (C) in proximal and distal dendrites of dSPNs (control, n = 10 cells from 6 mice; PD, n =11 cells from 5 mice; off-state, n = 13 cells from 8 mice; on-state, n = 10-11 cells from 7-8 mice).* p < 0.05, ** p< 0.01, **** p < 0.0001, Mann-Whitney test. (D) Confocal image of ChR2-YFP expression in the motor cortex in a coronal section. Scale bar is 1 mm. (E) Schematic diagram of the recording configuration. Whole-cell patch clamp recordings were made from dSPNs in acute brain slices of mice and oEPSCs were evoked by brief blue LED pulses. (F) Sample traces of Sr^2+^-oEPSC evoked by optogenetic stimulation of cortical afferents (indicated by blue vertical lines) in the presence of 3 mM Sr^2+^ and nominally 0 Ca^2+^ from dSPNs in LID off- and on-states. Detection and analysis of asynchronous release events was restricted to the window from 40 ms to 400 ms after stimulation. Scale bars denote 20 pA and 100 ms. (G) Cumulative probability plot of Sr^2+^-oEPSC amplitudes in dSPNs from off-state and on-state mice. (H) Box plot summary showing an increase in Sr^2+^-oEPSC amplitudes in on-state dSPNs (off-state, n = 10 cells from 4 mice; on-state, n = 8 cells from 4 mice). *** p < 0.001, Mann-Whitney test.

To determine whether these structural changes were accompanied by alterations in synaptic function, a combination of electrophysiological and optogenetic approaches was employed ^29,59,60^. To selectively activate corticostriatal synapses, an adeno-associated virus (AAV) carrying a channelrhodopsin 2 (ChR2) expression construct (AAV5-hSyn-hChR2(H134R)-EYFP) was injected into the motor cortex ipsilateral to the 6-OHDA lesion (Fig. 2D). In brain slices from off- and on-state *Drd1*-Tdtomato mice, dSPNs were patched with a Cs^+^-containing intracellular solution and voltage-clamped. To measure unitary synaptic strength, the extracellular Ca^2+^ was replaced with Sr^2+^ (3 mM), cortical fibers were stimulated with blue LED pulses and asynchronous, excitatory postsynaptic currents (Sr^2+^-oEPSCs) were recorded (Fig. 2, E-F). Consistent with the elevation in mushroom spine density, the distribution of Sr^2+^-oEPSC amplitudes was shifted toward larger values and the Sr^2+^-oEPSC amplitude was significantly larger in on-state dSPNs than in off-state dSPNs (p = 0.0003) (Fig. 2, F-H). Although the frequency of on-state Sr^2+^-oEPSCs trended toward higher values (Supplemental Fig. S1A), as with spine density (Fig. 2, B-C), the differences between states were not statistically significant (p = 0.127). Taken together, these observations suggest that on-state dopaminergic signaling in dyskinetic mice leads to an enhancement in intrinsic excitability and a potentiation of corticostriatal synaptic function in dSPNs, both of which are consistent with what is known about the consequences of D1R activation in dSPNs from healthy mice.

### Intrinsic excitability and synaptic connectivity of iSPNs decreased in the on-state of LID

Although it is clear that dSPNs are critical to the emergence of LID, iSPNs also are pivotal to this condition ^61,62^. Clear changes in the properties of iSPNs following LID induction have been reported, but previous assessments have been limited to the off-state, leaving unexplored how they are modulated in the on-state ^40–43^. To fill this gap, iSPNs were studied using *Drd2*-enhanced green fluorescent protein (eGFP) mice following unilateral 6-OHDA lesioning and levodopa treatment.

Using the same strategy to assess somatic excitability as described above (Fig. 3A), on-state iSPNs were found to be less excitable than in the off-state (Fig. 3, B-D): rheobase was elevated compared to the off-state (p = 0.0013) (Fig. 3D) and the F-I relationship curve shifted to the right (Fig. 3C).

**Figure 3.**
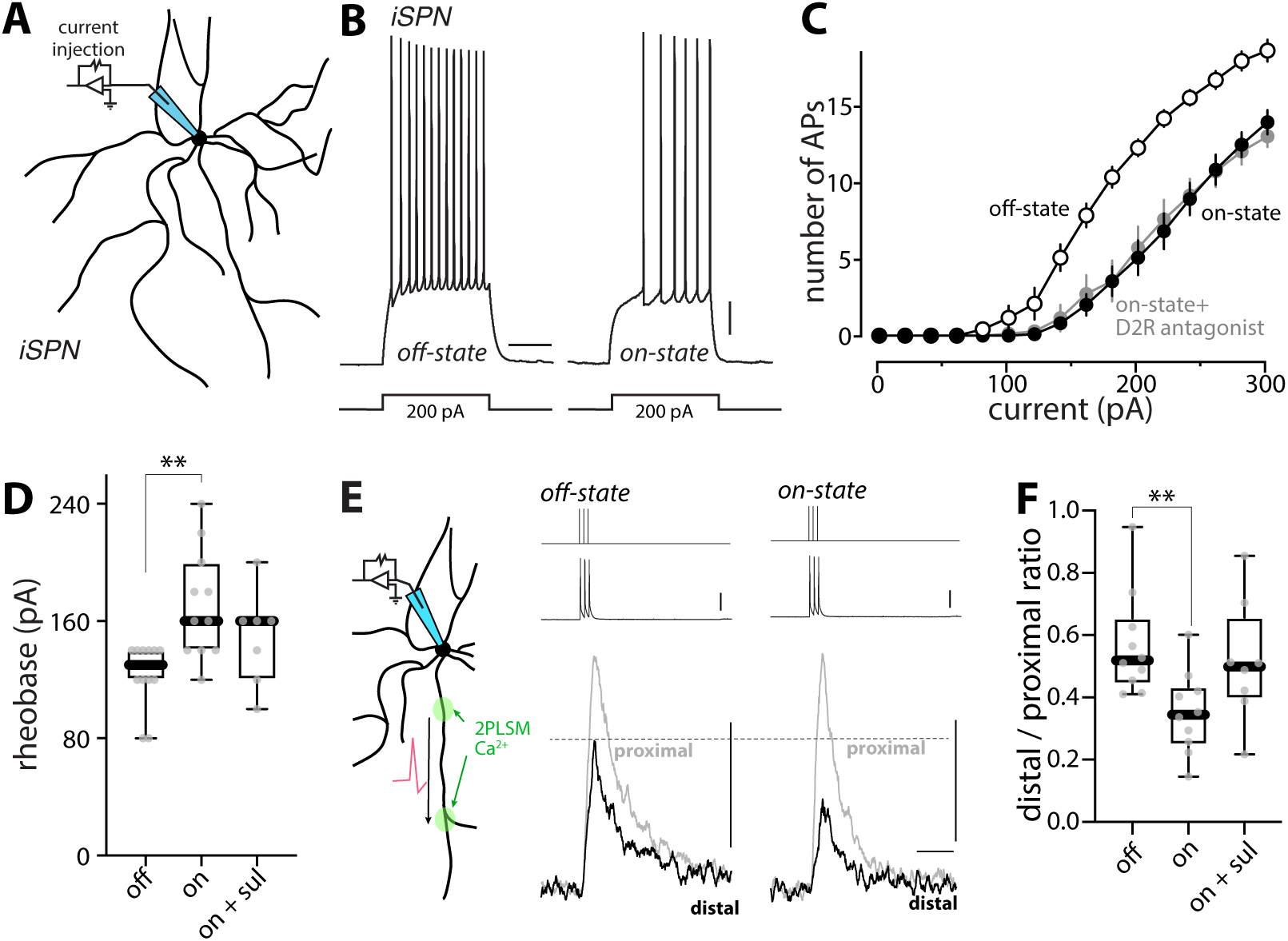
Somatic and dendritic excitability of iSPNs fluctuated between off- and on-states of LID. (A) Schematic illustrating the somatic excitability assay in iSPNs. (B) Sample voltage recordings from off-state and on-state iSPNs in response to a 200-pA current injection (500 ms duration). Scale bars are 10 mV and 200 ms. (C) Current-response curves showing the decrease in iSPN somatic excitability in the on-state which was resistant to bath application of D2R antagonist sulpiride (10 μM) (off-state, n = 12 cells from 5 mice; on-state, n = 11 cells from 8 mice; on-state + D2R antagonist, n = 7 cells from 4 mice). (D) Box plot summary of rheobase in iSPNs. ** p < 0.01, Mann-Whitney test. (E) Left, schematic illustrating the dendritic excitability assay in iSPNs. Right, Fluo-4 fluorescence transients (bottom) recorded from iSPN proximal and distal dendrites are aligned with current injections of 2 nA (top) and corresponding somatic voltage changes (middle). Scale bars denote 25 mV, 0.1 ΔG/R0 and 0.5 s. (F) Box plot summary of dendritic excitability index (the distal to proximal ratio of AUC of ΔG/R_0_) in iSPNs from off-state, on-state or on-state with bath application of sulpiride (off-state, n = 10 cells from 5 mice; on-state, n = 10 cells from 5 mice; on-state + sul, n = 8 cells from 6 mice). ** p < 0.01, Mann-Whitney test.

Acute antagonism of D2Rs with sulpiride (10 μM) in slices taken from on-state mice did not alter somatic excitability (rheobase, p = 0.287), arguing that the hypoexcitability was not being maintained by DA (Fig. 3, C-D). To complement the assessment of somatic excitability, the dendritic excitability of iSPNs also was examined using the combination of patch clamp electrophysiology and 2PLSM, as described above. In contrast to dSPNs, the index of dendritic excitability in iSPNs fell in the on-state (p = 0.0029) (Fig. 3, E-F). However, unlike the modulation of dSPN dendritic excitability, acute blockade of dopaminergic signaling did not significantly alter dendritic excitability in on-state iSPNs (p = 0.0545) (Fig. 3F), suggesting some other factor was involved.

As with D1Rs in dSPNs, D2R signaling in iSPNs modulates glutamatergic synaptic plasticity^9,10,55^. Activation of D2Rs promotes the induction of LTD at axospinous, corticostriatal synapses ^56,63^, which in principle should lead to a reduction in spine size ^64^. In addition, homeostatic mechanisms have been posited to regulate iSPN spine density ^24,65^. As a first step toward determining whether there were alterations in spine density or size in the on-state, iSPNs in *ex vivo* brain slices from *Drd2*-eGFP mice were examined after 6-OHDA lesioning and in the off-state after LID induction. In agreement with previous work ^24,40,42,65,66^, spine density in both proximal and distal dendrites was significantly reduced 3-4 weeks after a 6-OHDA lesion (p < 0.0001 in both proximal and distal dendritic locations) (Fig. 4, A-B). After LID induction, off-state iSPN spine density in both dendritic regions returned to a range that was indistinguishable from that of iSPNs from mice without 6-OHDA lesions (Fig. 4, A-B), as previously reported ^40,66^. Unexpectedly, in the on-state, the apparent spine density measured with 2PLSM fell significantly in in both proximal and distal iSPN dendrites (on-state vs. off-state: p < 0.0001 for both proximal and distal dendrites) (Fig. 4, A-B), suggesting that axospinous synapses were being added in the off-state and removed in the on-state. Similar changes were observed in the density of mushroom spines (on-state vs. off state: distal, p = 0.0006; proximal, p = 0.0002) (Fig. 4C).

**Figure 4.**
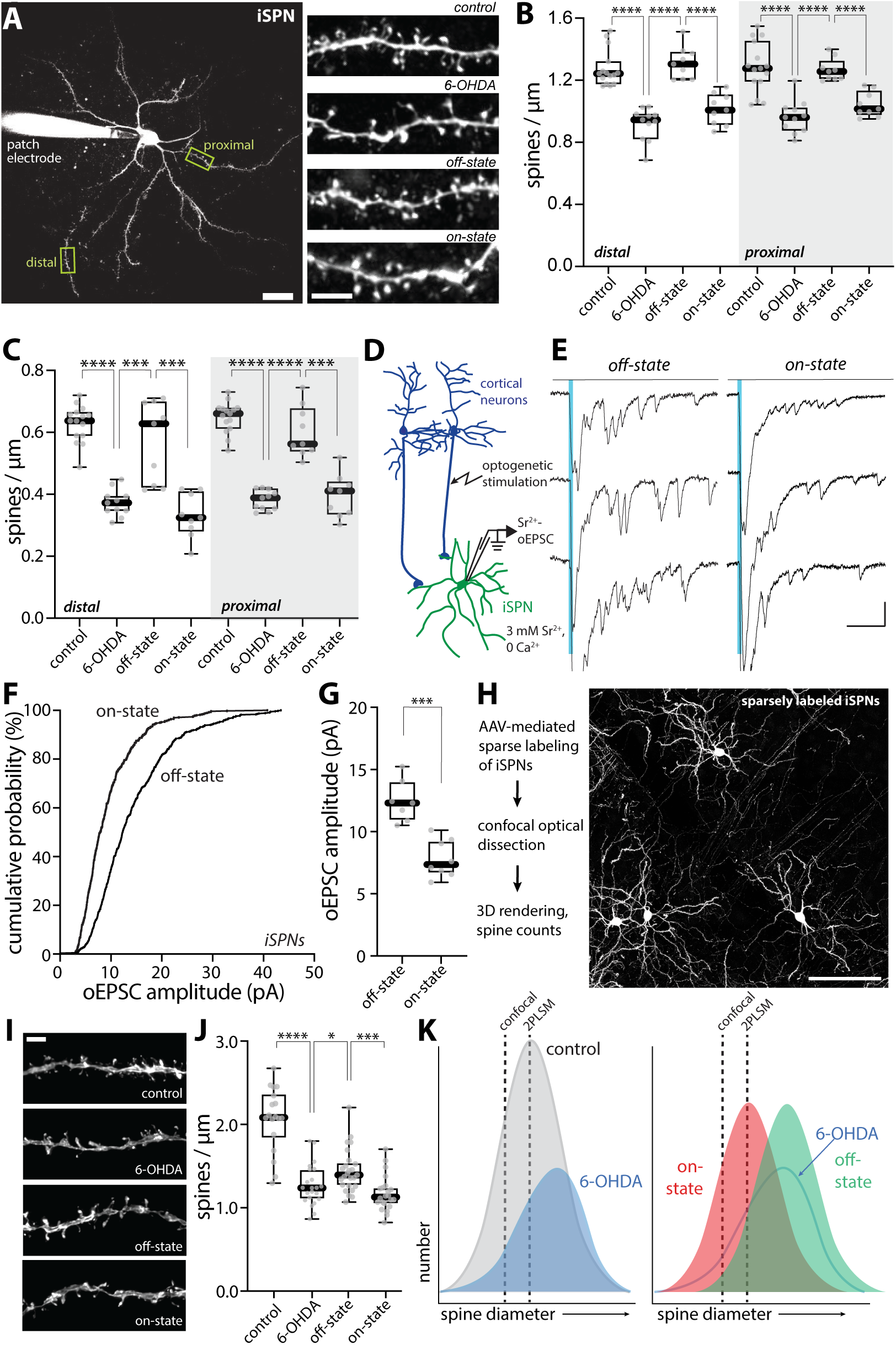
Spine morphology and unitary synaptic strength of iSPNs changed between LID-off and LID-on states. (A) Left, a low-magnification 2PLSM image of a patched iSPN with its dendritic tree visualized by Alexa dye and its proximal and distal dendritic segments delineated. Scale bar is 20 μm. Right, representative 2PLSM images of dendritic segments of iSPNs from control, 6-OHDA lesioned, LID off-state and LID on-state mice. Scale bar represents 5 μm. (B-C) Box plot summaries of total spine density (B) and mushroom spine density (C) in proximal and distal dendrites of iSPNs (control, n = 15 cells from 8 mice; PD, n = 11 cells from 6 mice; off-state, n = 9 cells from 5 mice; on-state, n = 9 cells from 5 mice). **** p < 0.0001, *** p < 0.001, Mann-Whitney test. (D) Schematic illustrating the experimental setup for Sr^2+^-oEPSC measurement. Whole-cell patch clamp recordings were made from iSPNs in acute brain slices of mice and oEPSCs were evoked by brief blue LED pulses in the presence of 3 mM Sr^2+^ and nominally 0 Ca^2+^. (E) Representative recordings of Sr^2+^-oEPSC evoked by optogenetic stimulation of cortical afferents (indicated by blue vertical lines) from iSPNs of off-state and on-state mice. Asynchronous release analysis was restricted to the window from 40 ms to 400 ms after stimulation. Scale bars denote 20 pA and 100 ms. (F) Cumulative probability plot of Sr^2+^-oEPSC amplitudes in iSPNs from off-state and on-state mice. (G) Box plot summary showing change in Sr^2+^-oEPSC amplitudes in iSPNs between LID off-and on-states (off-state, n = 7 cells from 4 mice; on-state, n = 8 cells from 4 mice). *** p < 0.001, Mann-Whitney test. (H) Left, workflow of confocal microscopy of sparsely labeled iSPNs. Right, high-resolution confocal image of sparsely labeled iSPNs in the DLS of a control mouse. Scale bar = 100 μm. (I) Representative images showing proximal dendritic segments of sparsely labeled iSPNs from control, 6-OHDA lesioned, LID off-state and LID on-state mice. Surrounding dendrites and axons were masked using the Surface function of Imaris. Scale bar, 3 μm. (J) Box plot summary of spine density in proximal dendrites of sparsely labeled iSPNs imaged by high-resolution confocal microscopy (control: n = 18 dendrites from 4 mice; 6-OHDA: n = 22 dendrites from 3 mice; off-state: n = 25 dendrites from 4 mice; on-state: n = 22 dendrites from 4 mice). **** p < 0.0001, *** p < 0.001, * p < 0.05, Mann-Whitney test. (K) Schematic showing different distributions of iSPN spine size in control, 6-OHDA lesioned, LID off-state and on-state mice, as well as the detection thresholds of 2PLSM and confocal microscopy. Note that the apparent changes in spine density between the on- and off-states revealed by 2PLSM was mostly attributable to changes in spine size (hence, detectability by 2PLSM imaging), rather than changes in spine population.

If there were oscillations in the number of axospinous synapses between on- and off-states, then there should be a corresponding change in the frequency of asynchronous oEPSCs generated by activation of the corticostriatal pathway when extracellular Ca^2+^ was replaced with Sr^2+^. To test this hypothesis, M1 cortical pyramidal neurons were induced to express ChR2 as described above and asynchronous, Sr^2+^-oEPSCs were recorded in iSPNs in *ex vivo* brain slices taken from mice in either the LID on- or off-states (Fig. 4, D-E). Unexpectedly, the amplitude, but not the frequency, of Sr^2+^-oEPSCs was significantly smaller in on-state iSPNs than in off-state iSPNs (amplitude: p = 0.0003; frequency: p = 0.23) (Fig. 4, F-G, also Supplemental Fig. S1B). These observations are consistent with the inference that what is changing in iSPNs between on- and off-states is the strength, but not the number, of corticostriatal axospinous synapses.

How can these seemingly disparate experimental results be reconciled? An obvious caveat to the 2PLSM counts is that thin spines can be missed. The density of SPN dendritic spines detected with high voltage electron microscopy (HVEM) is considerably greater than that estimated from three-dimensional 2PLSM reconstructions ^67,68^. Although HVEM was beyond our experimental reach, high-resolution confocal microscopy with fixed brain tissue (see Detailed Methods) does offer an alternative approach. To this end, *Adora2*-Cre mice were injected with a diluted AAV carrying a Cre-dependent eGFP expression construct (AAV9-pCAG-flex-EGFP-WPRE). This led to sparse labeling of iSPNs, allowing visualization of individual dendrites (Fig. 4, H-I). Three-dimensional confocal reconstruction of iSPNs yielded spine density estimates in proximal dendrites that were significantly higher than those estimated from 2PLSM optical sectioning of live tissue (Fig. 4J) (confocal microscopy: median 2.084 vs. 2PLSM: median 1.243, spines per μm). After 6-OHDA lesioning, iSPN spine density estimated from confocal imaging decreased (control vs. PD: p < 0.0001), consistent with previous reports ^24,40,42,65,66^. However, in contrast to the conclusions drawn from 2PLSM, confocal spine density estimates did not return to control values in the off-state after LID induction, although they did increase modestly from the 6-OHDA lesioned state (p = 0.0136) (Fig. 4, I-J). As with 2PLSM, confocal iSPN spine density estimates fell in the on-state (p = 0.0001) (Fig. 4, I-J), but only modestly.

What these observations suggest is that the apparent fluctuations in iSPN spine density following LID induction largely reflect alterations in the size – and detectability – but not the number of axospinous synapses. To illustrate how this might occur, the distribution of spine diameters was hypothesized to Gaussian (Fig. 4, K). Given the limitations of light microscopy, there is a threshold diameter for spine detection; this threshold should be greater for 2PLSM which used a lens with relatively smaller numerical aperture (NA) than that used for the confocal analysis (0.9 NA 60X lens for 2PLSM, 1.49 NA 60X lens for confocal microscopy); hypothetical thresholds are drawn on the distributions to show how in principle they should change spine density estimates (Fig. 4K). After DA depletion, the distribution of spine diameters should change in two ways. First, the total number of synapses should drop by about 30% to reflect frank spine pruning. Second, the distribution of diameters should shift slightly toward larger values to account for the aberrant potentiation of axospinous synapses accompanying loss of iSPN D2R signaling ^40^ (Fig. 4K). To illustrate the alterations between on- and off-states, the distribution of spine diameters was simply shifted to the right (larger) to reflect synaptic potentiation in the off-state and to the left (smaller) in the on-state to reflect synaptic depression in the on-state (Fig. 4K). This simple model accounts for the *apparent* alterations in spine density estimated with 2PLSM, as well as the Sr^2+^-oEPSC data.

### ACh release was elevated in PD and LID off-state and inhibited by D2R signaling

The data presented to this point argue that dyskinesiogenic fluctuations in dopaminergic signaling between the on- and off-states drive cell-specific changes in the intrinsic excitability and functional connectivity of iSPNs and dSPNs. Fluctuations in brain DA trigger these changes, but it is unclear whether, at the level of individual SPNs, DA is acting alone. That is, are there other alterations in the striatal circuitry induced by levodopa treatment that contribute to the observed changes in the functional properties of SPNs? One key target of intrastriatal dopaminergic signaling is the ChI. Autonomously active ChIs are robustly inhibited by DA acting on D2Rs ^15–18,69^. Moreover, ACh potently modulates the intrinsic excitability and synaptic plasticity of iSPNs and dSPNs through mechanisms that complement those of DA ^70,71^. However, the role of ChIs in the parkinsonian state and LID is controversial (reviewed by ^26,34–36^).

To directly assess ACh release by ChIs in the parkinsonian and LID states, the genetically encoded ACh sensor GRAB_ACh3.0_ was used in conjunction with 2PLSM ^72^. About three weeks after stereotaxic injection into the dorsolateral striatum (DLS) of an AAV carrying a GRAB_ACh3.0_ expression construct, *ex vivo* brain slices were prepared and sensor fluorescence monitored in response to local electrical stimulation (Fig. 5, A-B). Robust ACh signals were evoked by either single pulses (Fig. 5) or short pulse bursts (Supplemental Fig. S2). In the healthy DLS, the ACh signal rose rapidly and decayed back to baseline within 1-2 seconds (Fig. 5C). Bath application of the D2R antagonist sulpiride (10 µM) increased the evoked ACh signal (p = 0.0005) (Fig. 5, C, F; Supplemental Fig. S2), indicating that there was a basal dopaminergic tone. As expected, bath application of a D2R agonist quinpirole (10 µM) suppressed the ACh signal (p = 0.0034) (Fig. 5, C, F; Supplemental Fig. S2). In *ex vivo* slices from 6-OHDA lesioned mice, the GRAB_ACh3.0_ signal was dramatically elevated (unlesioned vs. 6-OHDA: p < 0.0001) (Fig. 5, B,D,F; Supplemental Fig. S2), suggesting that ACh release was disinhibited by DA depletion. Consistent with this scenario, bath application of sulpiride (10 μM) had no effect (p = 0.5566), whereas D2R agonists (50 nM DA or 10 μM quinpirole) strongly inhibited ACh release in 6-OHDA mice (DA: p = 0.002; quinpirole: p = 0.002) (Fig. 5, D,F; Supplemental Fig. S2). Interestingly, the peak GRABACh3.0 fluorescence in 6-OHDA-lesioned DLS was significantly greater than that in the control DLS in the presence of the D2R antagonist (p < 0.0001), consistent with previous work suggesting that muscarinic autoreceptor function in ChIs was impaired following DA depletion ^27^. In slices from LID off-state mice, ACh release was significantly greater than that in the unlesioned striatum (p < 0.0001), but similar to that following a 6-OHDA lesion (p = 0.4679). Bath application of DA (50 nM), at a concentration comparable to that detected in the striatum *in vivo* during on-state ^73–75^, strongly suppressed ACh release (p = 0.001) (Fig. 5, E-F; Supplemental Fig. S2). These data suggest that ACh release falls in the on-state as intrastriatal DA levels rise and then rebounds in the off-state when DA levels drop.

**Figure 5:**
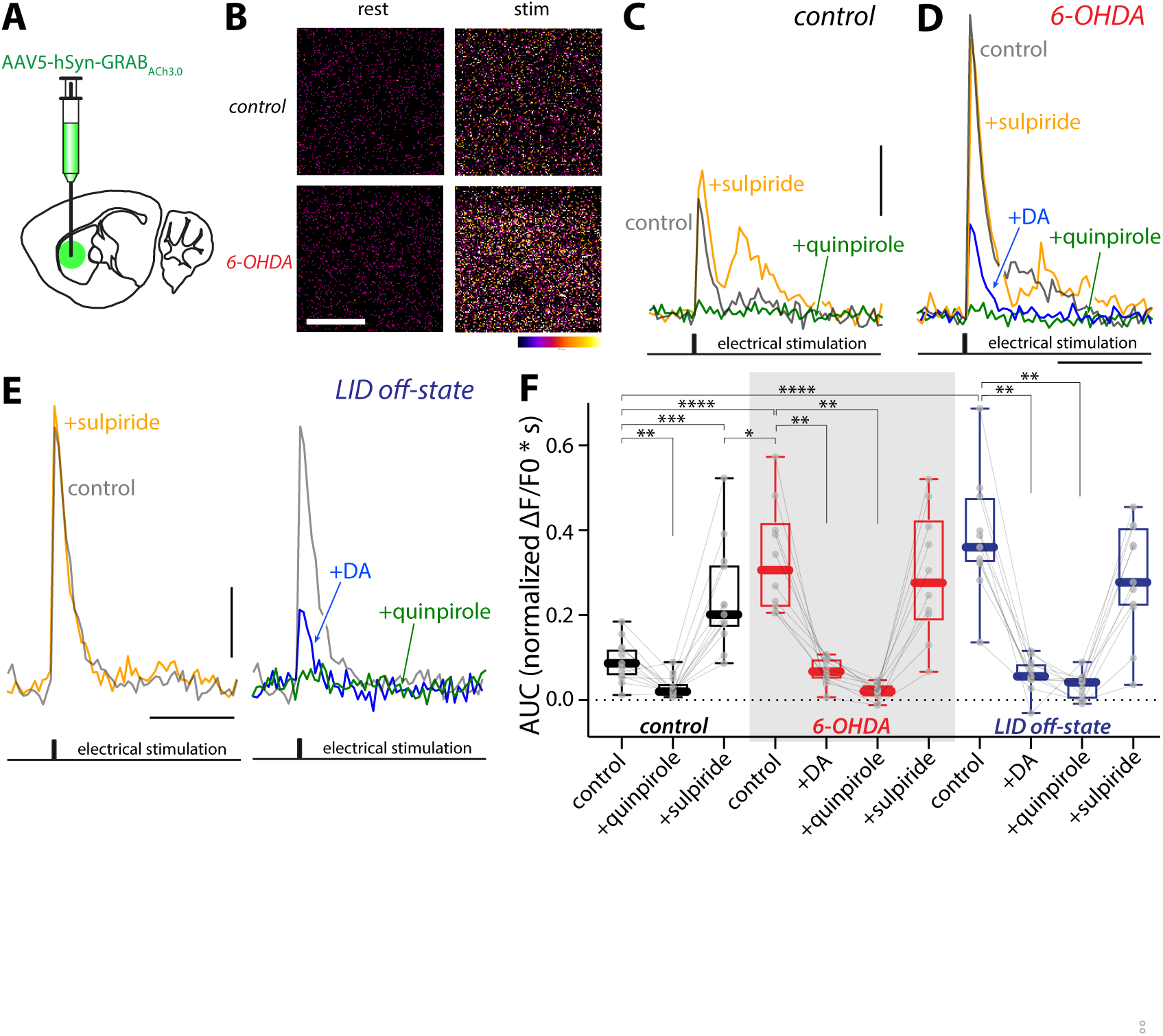
ACh release was increased in the striata of PD and LID mice but strongly suppressed by DA. (A) Diagram showing injection of GRAB_ACh3.0_-expressing AAV into the DLS of mice. (B) Pseudocolor images of GRAB_ACh3.0_ signal (ΔF/F_0_) in resting state (‘rest’) and immediately following stimulation (‘stim’) in the DLS of unlesioned (‘control’) or 6-OHDA lesioned (‘6-OHDA’) mice. Scale bar is 20 μm. (C-E) Representative traces of the fluorescence response of GRAB_ACh3.0_ signals in the DLS of control (C), 6-OHDA lesioned (D), and LID off-state mice (E) evoked by an intrastriatal electrical stimulation before and after bath application of D2R agonist(s) (quinpirole or DA) or D2R antagonist sulpiride. Scale bars are 1 s and 0.2 ΔF/F_0_. (F) Box plot summary of GRAB_ACh3.0_ signal (AUC of normalized ΔF/F_0_) in unlesioned control, 6-OHDA lesioned, and LID off-state mice. In 6-OHDA lesioned and off-state mice, ACh release was significantly elevated. Bath application of DA (50 nM), mimicking the high DA condition in LID on-state, strongly suppressed ACh release (unlesioned, n = 13 ROIs from 3 mice; 6-OHDA, n = 10 ROIs from 3 mice; off-state, n = 11 ROIs from 5 mice). * p < 0.05, ** p < 0.01, *** p < 0.001, **** p < 0.0001, Mann-Whitney (unpaired) and Wilcoxon (paired) tests.

### M1 muscarinic receptor-CDGI signaling contributed to iSPN plasticity and LID induction

How might the dysregulation of ACh release have contributed to the changes in SPN properties found in LID on- and off-states? The intrinsic excitability and synaptic function of both iSPNs and dSPNs are potently modulated by muscarinic acetylcholine receptors (mAChRs). Previous work has implicated an imbalance in the activity of dSPN D1Rs and M4Rs during the on-state in LID induction ^20,37^. How alterations in iSPN properties were influenced by disrupted cholinergic signaling is less clear-cut. In normal mice, activation of iSPN M1Rs enhances both somatic and dendritic excitability ^22–24^. Genetic deletion of the gene coding for CDGI, which couples M1Rs to intracellular signaling pathways, blunts the M1R-induced modulation of iSPNs dendritic excitability and prevents the induction of LTP at corticostriatal glutamatergic synapses ^22^.

To assess the role of M1Rs in LID-induced adaptations in iSPNs, dyskinetic *Drd2* BAC mice were given intraperitoneal injections of the M1R antagonist trihexyphenidyl hydrochloride (THP, 3 mg/kg) or saline at the beginning of the off-state (∼2 hr after levodopa administration); THP is known to have good brain bioavailability and favorable pharmacokinetics after peripheral administration ^76^.

Mice were sacrificed 16 hours later, *ex vivo* brain slices prepared and iSPNs patch clamped (Fig. 6A). The slices from mice given THP were continuously incubated with the M1R selective antagonist VU 0255035 (5 µM) throughout the experiment. Somewhat unexpectedly, M1R antagonism did not have a significant impact on the elevation in iSPN somatic excitability in the off-state (Fig. 6, B-C). However, M1R antagonism did prevent the off-state elevation in dendritic excitability (p = 0.012) (Fig. 6D) and the apparent elevation in spine density (p = 0.0003) (Fig. 6, E-F).

**Figure 6.**
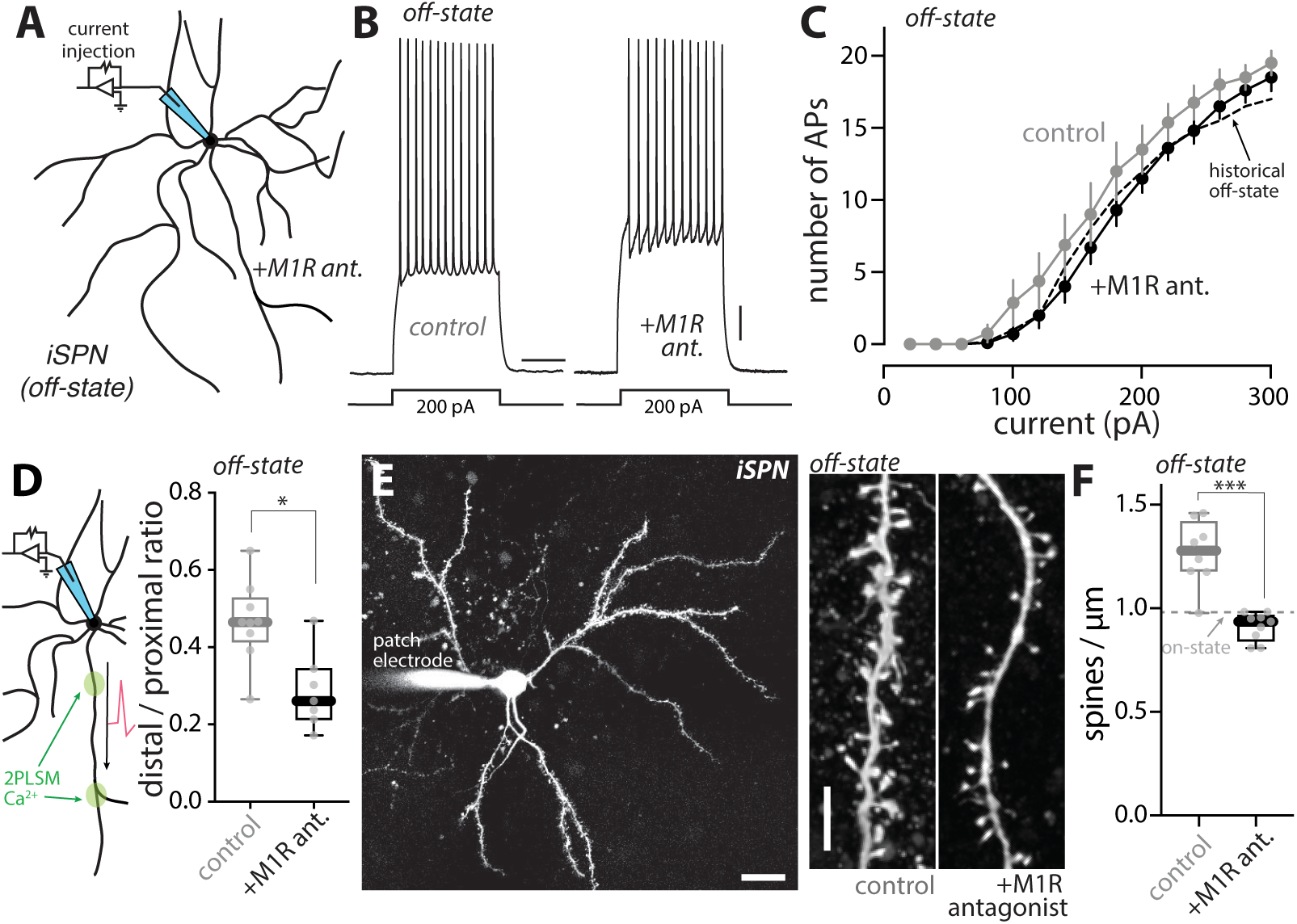
M1R mediated the dendritic, but not the somatic, alterations in iSPNs in LID off-state. (A) Schematic illustrating the somatic excitability assay in iSPNs from LID off-state mice in which M1R signaling was inhibited throughout the off-state. (B) Sample voltage changes evoked by 200-pA current injections (500-ms duration) in iSPNs from off-state mice that received M1R antagonist or vehicle control throughout the off-state. Scale bars are 10 mV and 200 ms. (C) Current-response curves showing that iSPN somatic excitability in off-state was resistant to M1R antagonists (off-state control, n = 8 cells from 4 mice; off-state with M1R antagonists, n = 10 cells from 5 mice). Historical off-state data from Fig. 3C was overlaid as a dashed line. (D) Left, schematic illustrating the dendritic excitability assay in iSPNs. Right, summary of dendritic excitability index in iSPNs from off-state mice treated with M1R antagonist or vehicle control (off-state control, n = 9 cells from 5 mice; off-state+M1R antagonists, n = 7 cells from 4 mice). * p < 0.05, Mann-Whitney test. (E) Left, low-magnification image showing an iSPN from an off-state mouse treated with M1R antagonists, patched and filled with Alexa dye. Scale bar is 20 μm. Right, sample 2PLSM images of dendritic segments of iSPNs from off-state mice treated with vehicle (‘control’) or M1R antagonist (‘+M1R antagonist’). Scale bar is 5 μm. (F) Box plot summary of dendritic spine density in off-state iSPNs treated with vehicle or M1R antagonists (off-state control, n = 8 cells from 4 mice; off-state+M1R antagonist, n = 9 cells from 5 mice). *** p < 0.001, Mann Whitney test. Historical on-state data from Fig. 4B was shown by a dashed line.

These observations suggest that M1R signaling contributes to the LID-induced adaptations in iSPNs, particularly those in dendrites. The obvious limitations of these experiments are that THP is not selective for M1Rs ^77^ and that M1Rs are broadly distributed in the brain – raising the possibility that the effects of THP are not dependent upon iSPN M1Rs. As a first step toward addressing these limitations, CDGI knockout (KO) mice were studied; CDGI is largely restricted to the striatum and is a key mediator of dendritic M1R signaling in iSPNs ^22,25^. CDGI KO mice were subjected to the same LID induction protocol as described above and then brain slices prepared for study. In agreement with its dendritic role ^22^, deletion of CDGI did not blunt the oscillation in iSPN somatic excitability between LID on- and off-states (Fig. 7, A-C). To assess dendritic excitability, bAP propagation was examined. In *ex vivo* brain slices from naïve wildtype mice, bath application of the mAChR agonist oxo-M (10 µM) robustly increased the invasion of bAPs into the dendrites of iSPNs (p = 0.0098) (Fig. 7, D-E), in agreement with previous work ^23^. However, in *ex vivo* slices from naïve mice lacking CDGI, the M1R agonist oxo-M had no effect on dendritic invasion of bAPs (p = 0.910) (Fig. 7, E). Consistent with this observation, the enhancement of bAP dendritic invasion in wildtype iSPNs in the LID off-state was not seen in off-state CDGI KO iSPNs (off-state vs. on-state: wildtype, p = 0.0029; CDGI KO, p = 0.395) (Fig. 7F). Furthermore, the apparent increase in iSPN spine density observed in wildtype mice in the off-state (Fig. 4B) was not seen in iSPNs from CDGI KO mice (off-state vs. on-state: CDGI KO, p = 0.613) (Fig. 7, G-H). Taken together, these observations suggest that dendritic M1R/CDGI signaling in iSPNs plays a key role in the structural and functional adaptations accompanying the off-state in dyskinetic mice.

**Figure 7:**
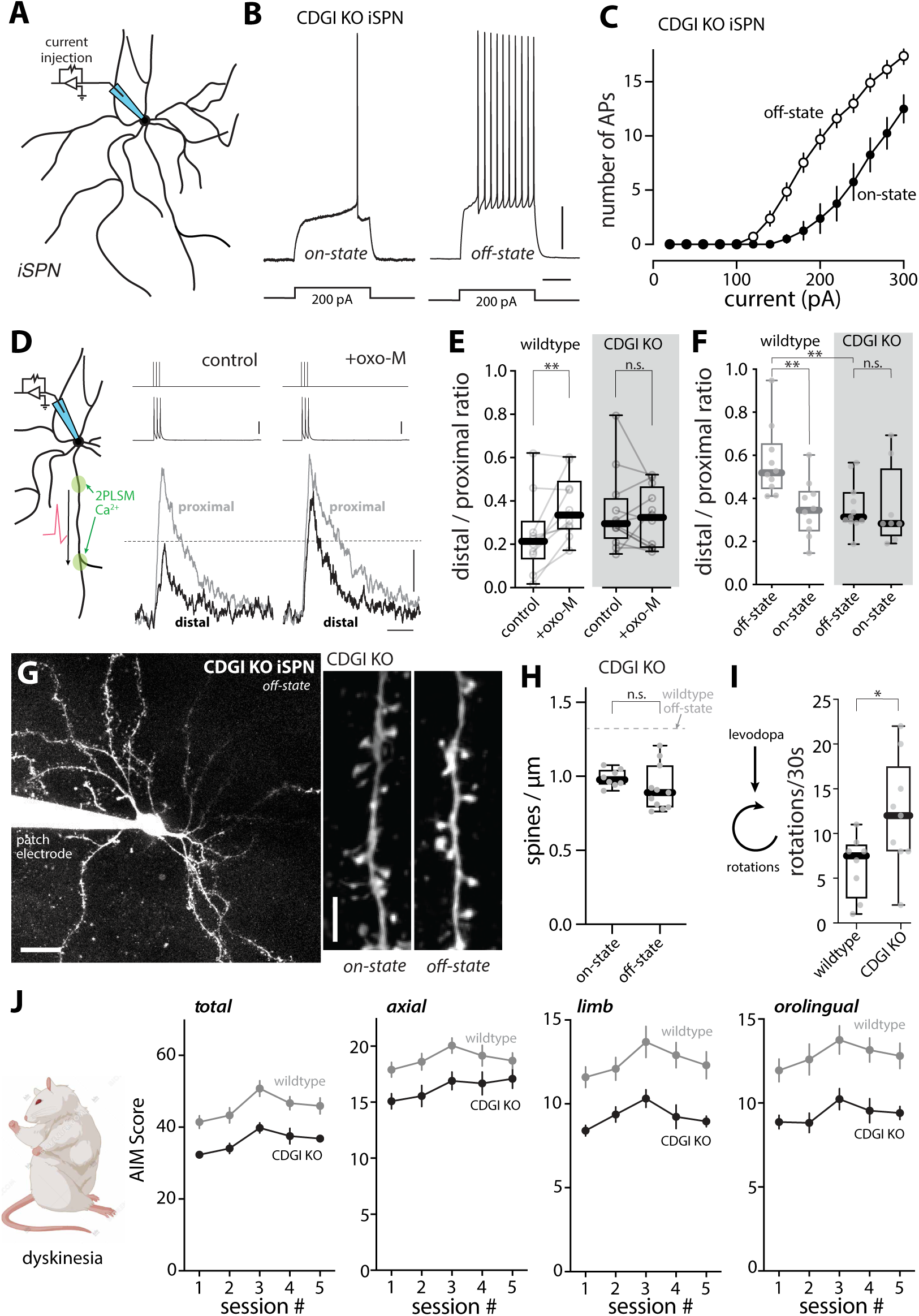
CDGI signaling pathway mediated the dendritic changes in off-state iSPNs and contributed to dyskinetic behavior. (A) Schematic illustrating the somatic excitability assay in iSPNs from CDGI KO mice. (B) Sample voltage recordings from on-state and off-state iSPNs from CDGI KO mice in response to a 200-pA current injection (500-ms duration). Scale bars are 10 mV and 200 ms. (C) Current-response curves showing that the fluctuation in iSPN somatic excitability between on- and off-states was not prevented by genetic deletion of CDGI (on-state, n = 8 cells from 4 mice; off-state, n = 13 cells from 5 mice). (D) Left, schematic illustrating the dendritic excitability assay in iSPNs with oxo-M application. Right, fluorescence transients of Fluo-4 from iSPN proximal and distal dendrites before and after application of oxo-M (10 μM) in slices from wildtype mice. Scale bars denote 50% ΔF/F_0_ and 0.5 s. Current injections of 2 nA and corresponding somatic voltage changes were shown on top and temporally aligned (scale bar denotes 25 mV). (E) Box plot summary showing the effect of oxo-M on iSPN dendritic excitability in wildtype and CDGI KO mice (wildtype, n = 10 dendrites from 5 cells/3 mice; CDGI KO, n = 12 dendrites from 6 cells/3 mice). ** p < 0.01, n.s., no significance, Wilcoxon test. (F) Box plot summary of dendritic excitability index of iSPNs from wildtype or CDGI KO mice that were in off- or on-state (wildtype data were from Fig. 3F; wildtype off-state, n = 10 cells from 5 mice; wildtype on-state, n = 10 cells from 5 mice; CDGI KO off-state, n = 11 cells from 5 mice; CDGI KO on-state, n = 9 cells from 5 mice). ** p < 0.05, n.s., no significance, Mann-Whitney test. (G) Left, low-magnification image of an iSPN from an off-state CDGI KO that was filled with Alexa dye through the patch pipette. Scale bar is 20 μm. Right, sample images of iSPN dendrites from CDGI KO mice in on- or off-state of LID. Scale bar is 5 μm. (H) Box plot summary of iSPN spine density in on- and off-state CDGI KO mice (on-state CDGI KO, n = 8 cells from 4 mice; off-state CDGI KO, n = 11 cells from 6 mice). n.s. not statistically significant, Mann-Whitney test. Historical off-state data from wildtype mice (Fig. 4B) was indicated by a dashed line. (I) Box plot summary of the number of contralateral rotations (in 30 seconds) recorded 40 min after the fifth levodopa administration in wildtype or CDGI KO mice (wildtype n = 8 animals; CDGI KO, n = 9 animals). * p < 0.05, Mann-Whitney test. (J) Plots of total, axial, limb and orolingual AIM scores as a function of sessions in wildtype and CDGI KO mice (wildtype n = 10 mice; CDGI KO, n = 11 mice). Data are mean ± SEM.

To determine whether these CDGI-dependent adaptations were linked to behavior, the dyskinesia induced in wildtype and KO mice was compared using the protocol described above. CDGI KO and wildtype controls were subjected to a unilateral 6-OHDA lesion and about a month later evaluated for the extent of the lesion using the cylinder test. Mice with a near-complete lesion were then injected with dyskinesiogenic doses of levodopa every other day and their abnormal involuntary movements (AIMs) were rated using an established rating scale ^4,20,40^. Routinely, the pro-kinetic effects of levodopa were assessed by monitoring the number of contralateral rotations in a 30 second period during peak-dose dyskinesia ^78^. Interestingly, the pro-kinetic effects of levodopa were initially similar in wildtype and CDGI KO mice, but, by the fifth dose, the effects of levodopa on rotations were significantly greater in CDGI KO mice (Fig. 7I; Supplemental Fig. S3A). More importantly, axial, limb and orolingual AIMs, as well as total AIMs, were less severe in CDGI KO mice than wildtype controls (Fig. 7J). These results suggest that CDGI-dependent M1R signaling in iSPN dendrites makes an important contribution to the network pathophysiology underlying LID.

Although these results clearly implicate iSPNs in LID, it is possible that the loss of CDGI in other cell types (e.g., dSPNs) was a factor in the attenuation of LID. To address this caveat, a CRISPR-Cas9 approach was used to disrupt the expression of M1R specifically in iSPNs. In short, a mixture of two AAVs were injected into six sites across the ipsilateral DLS of unilaterally 6-OHDA lesioned *Adora2*-Cre mice that had passed the cylinder test (Fig. 8A); one of the AAVs carried a Cre-dependent Cas9 expression construct and the other carried *Chrm1*-targeting guide ribonucleic acid sequences (gRNAs) with a FusionRed (FR) reporter construct. Control mice were injected with only the gRNA-FR vector (Fig. 8A). In a subset of experiments in which *ex vivo* analysis was to be performed, a third AAV carrying a Cre-dependent enhanced yellow fluorescent protein (eYFP) reporter (AAV9-EF1a-DIO-EYFP) was added. Roughly a month later, mice were subjected to the LID induction protocol (Fig. 8A). The AAVs expressed well (Fig. 8B) and the ability of the mAChR agonist oxo-M to enhance somatic excitability of iSPNs patch clamped in *ex vivo* brain slices was consistently lost (Supplemental Fig. S4), demonstrating the efficacy of M1R deletion by the CRISPR-Cas9 approach.

**Figure 8.**
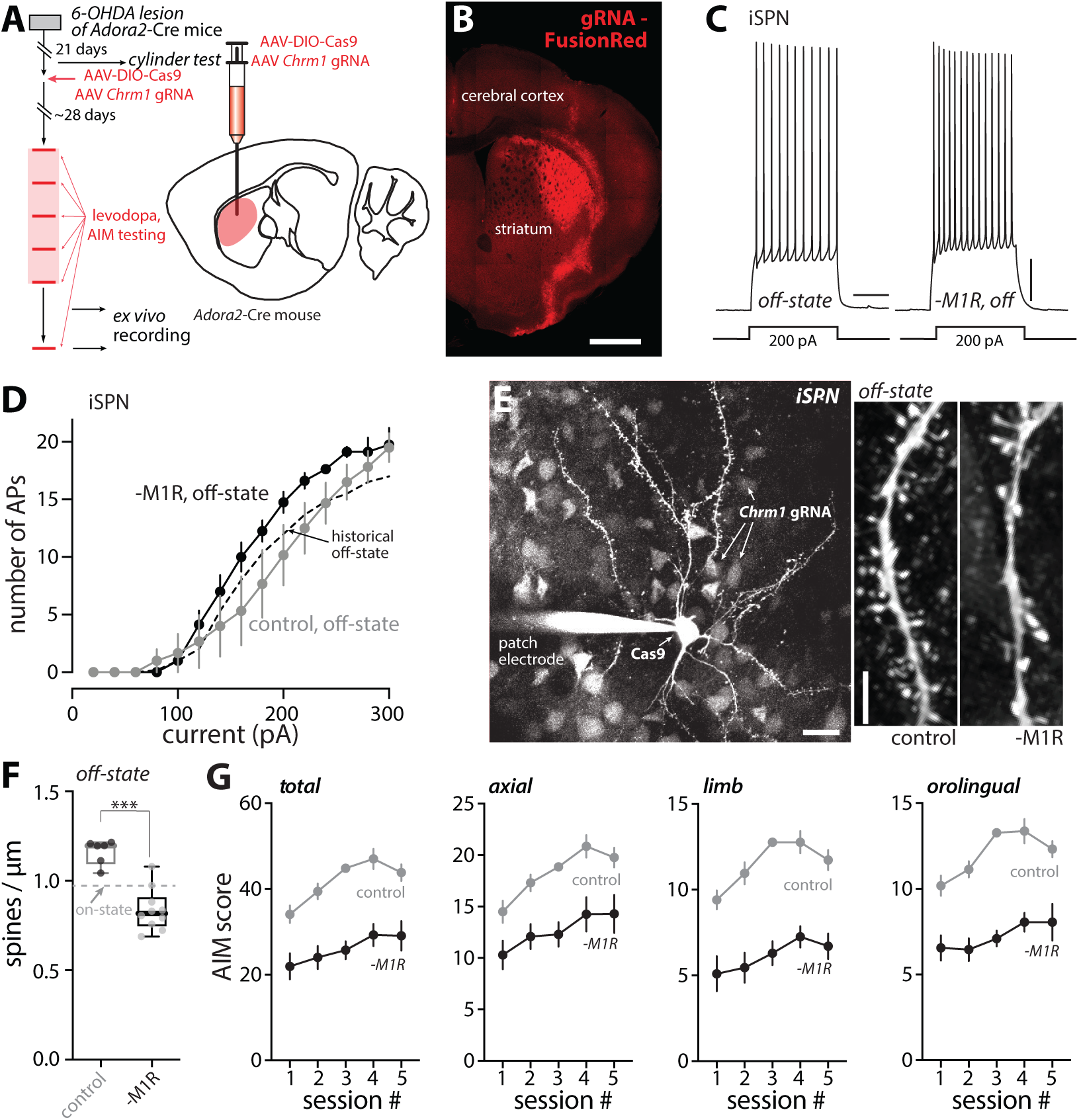
Deletion of M1R from iSPNs prevented dendritic changes in the off-state and attenuates dyskinetic behaviors. (A) Left, experimental timeline for 6-OHDA lesioning, M1R CRISPR expression, AIM testing and *ex vivo* recordings. Right, schematic illustrating injection of M1R CRISPR into the DLS of *Adora2*- Cre mice. (B) Confocal image showing expression of gRNA-FusionRed in the DLS of a coronal section. Scale bar is 1mm. (C) Sample somatic voltage changes in response to 200-pA current injections in iSPNs from off-state mice without or with M1R deletion from iSPNs. Scale bars are 10 mV and 200 ms. (D) Current-response curves showing that the increase in iSPN somatic excitability in the off-state was not prevented by M1R CRISPR (control: n = 6 cells from 3 mice; M1R deletion: n = 8 cells from 4 mice). Historical off-state data from Fig. 3C was shown by a dashed line. (E) Left, low-magnification image showing a patched iSPN with its dendritic arbor visualized by Alexa dye and expression of gRNA-FusionRed in nearby cells in a non-specific manner. Cell type-specific deletion was achieved by Cre-dependent expression of Cas9 in *Adora2*-Cre mice. Scale bar is 20 μm. Right, 2PLSM images of dendritic segments of iSPNs from off-state mice, without or with M1R deletion. Scale bar indicates 5 μm (F) Box plot summary of dendritic spine density of iSPNs without or with M1R deletion from off-state mice (control: n = 6 cells from 3 mice; M1R deletion: n = 10 cells from 4 mice). *** p < 0.001, Mann-Whitney test. Historical on-state data from Fig. 4B was indicated by a dashed line. (G) Plots of total, axial, limb and orolingual AIM scores as a function of sessions (data are mean ± SEM) in mice without or with iSPN-specific deletion of M1R. Genetic deletion of M1R in iSPNs produced an overall reduction in AIM scores (control n = 11 animals; M1R CRISPR, n = 10 animals).

To determine the cellular consequences of M1R deletion after LID induction, visually identified iSPNs in *ex vivo* brain slices were studied with patch clamp and 2PLSM approaches. As predicted from the systemic administration of the M1R antagonist (Fig. 6, B-C), genetic deletion of M1Rs from iSPNs did not alter the enhancement of somatic excitability observed in the off-state after LID induction (Fig. 8, C-D). Because the emission spectrum of FR and eYFP overlapped with those of the Ca^2+^ dyes, the bAP invasion experiments described above could not be performed.

Nevertheless, the dendritic architecture of iSPNs was examined by 2PLSM optical sectioning (Fig. 8, E-F). In agreement with the experiments employing systemic M1R antagonists or genetic deletion of CDGI, the apparent increase in iSPN spine density in the off-state following LID induction was blunted by genetic deletion of M1Rs (p = 0.0005) (Fig. 8, E-F). Behavioral analysis of mice in which M1Rs had been deleted selectively from iSPNs yielded results that were similar to those obtained from CDGI KO mice. First, the therapeutic benefit of levodopa (as assessed by contralateral rotations) rose as dyskinesia developed in mice lacking M1Rs in iSPNs (Supplemental Fig. S3B).

Second, axial, limb and orolingual AIMs were dramatically attenuated by deletion of M1Rs in iSPNs (Fig. 8G). These results, and those described above, argue that dendritic M1R/CDGI signaling in iSPNs contributes to structural and functional adaptations that cause the striatal pathophysiology underlying LID.

## Discussion

There are three main conclusions that can be drawn from the data presented. First, in both dSPNs and iSPNs, there was a profound oscillation in both intrinsic excitability and synaptic connectivity between on- and off-states following induction of LID in a mouse model of PD. These cellular and circuitry changes were complementary in nature, reinforcing the idea that an imbalance in the activity of dSPN and iSPN ensembles contributed to LID severity. Second, in parkinsonian and off-state dyskinetic mice there was a dramatic elevation in evoked ACh release, reflecting the loss of inhibitory D2R signaling. However, D2R signaling continued to robustly inhibit ACh release in tissue from dyskinetic mice, arguing that the oscillation in striatal DA concentrations associated with LID on- and off-states was mirrored by a counter-oscillation in ACh concentration. Third, the impact of cholinergic signaling on LID-associated adaptations in SPNs was particularly prominent in iSPNs. Indeed, blunting M1R or CDGI signaling in iSPNs not only attenuated the oscillations in dendritic excitability and synaptic strength, it also had clear behavioral effects – increasing the symptomatic benefit of levodopa and attenuating on-state dyskinesia.

### LID was associated with cell- and state-specific adaptations, particularly in dendrites

Although previous studies have shown that LID induction is accompanied by physiological and anatomical changes in SPNs, most of the studies have focused on alterations that are evident in the off-state, long after the last levodopa dose ^40–43^. An implicit assumption of many of these studies is that there are unlikely to be major changes in the relatively short period (hours) after levodopa treatment. Our studies show that this assumption is incorrect.

In dSPNs recorded in *ex vivo* brain slices taken from mice shortly after their last dose of levodopa (on-state), somatic and dendritic excitability was significantly greater than in dSPNs recorded in slices taken from off-state mice. This observation is not surprising given the ability of DA acting at D1Rs to increase the excitability of dSPNs at membrane potentials near spike threshold ^47–50^. Interestingly, the augmentation in somatic excitability did not readily reverse with antagonism of D1Rs, in keeping with the observation that in perforated patch recordings, this modulation is relatively persistent ^53^. In contrast, the elevation in dendritic excitability was more time-locked to D1R signaling, as it was acutely reversed by an antagonist. Why there is this difference in offset kinetics is unclear at this point but it is likely to involve the regulation of protein phosphatases that reverse the covalent modifications to proteins achieved by D1R activation of protein kinase A (PKA) ^49^. Accompanying these changes in excitability were state-dependent alterations in synaptic function. As previously reported, the density of 2PLSM-visible spines on dSPN dendrites fell with LID induction and termination of levodopa ^40^. The mechanisms driving this synaptic change are unclear. What is known is that if levodopa treatment is withheld for several months after 6-OHDA lesioning, 2PLSM estimates of dSPN spine density eventually fall ^58^. Given that ChIs are disinhibited in this condition, this shift could reflect the sustained engagement of M4R-dependent LTD. The ability of intermittent levodopa treatment to precipitate a similar synaptic attenuation could reflect network activation of disinhibited ChIs and more robust LTD induction ^20,79^. Regardless, right after levodopa treatment, the density of detectable spines, particularly those with a mushroom morphology, rose in dSPNs – as might be predicted by engagement of D1Rs and the induction of LTP at axospinous, glutamatergic synapses ^56,57^. Consistent with this inference, the amplitude of cortically evoked Sr^2+^-oEPSCs rose during the on-state. Both of these measures of synaptic strength reversed in the off-state, suggesting that the loss of dopaminergic signaling led to depotentiation or LTD ^20,43^. Importantly, the frequency of Sr^2+^-oEPSCs did not change between states, suggesting that there was not a significant change in the number of synapses between on- and off-states. From a network perspective, it is important to note that this drug-induced oscillation in synaptic strength was not driven by the usual factors governing DA release and engagement of D1Rs, like action initiation or reward. This observation suggests that there is a progressive randomization of corticostriatal synaptic strength with repeated levodopa treatment. If information that enables striatal dSPN ensembles to promote contextually appropriate action is stored in the strength of corticostriatal synapses, then the induction of LID should significantly degrade this information and the ability to move in a purposeful manner ^80,81^.

As with dSPNs, the properties of iSPNs in dyskinetic mice shifted between on- and off-states. In the on-state, somatic and dendritic excitability of iSPNs was lower than in the off-state. Unexpectedly, the suppression of iSPN excitability in the on-state did not readily reverse with D2R antagonism. The mechanisms responsible for this persistence, as well as its eventual reversal in the off-state, are unclear. One possibility is that on-state D2R signaling produces long-lasting alterations in ion channels governing excitability (i.e. intrinsic plasticity ^82,83^). In iSPNs, D2Rs could bring about this modulation through activation of phospholipase C and the protein phosphatase calcineurin, which regulates phosphorylation of a variety of ion channels, including Ca_v_1 Ca^2+^ channels ^84^. In contrast, the reasons for the elevation in off-state excitability of iSPNs are easier to surmise. Loss of ambient D2R signaling and disinhibition of ChIs undoubtedly contributed to this shift ^22–24,85^. Notably, pharmacological antagonism of M1Rs or genetic deletion of M1Rs/CDGI, blunted the off-state shift in dendritic excitability.

Another state-dependent change in iSPNs was in the strength of corticostriatal glutamatergic synapses. The amplitude of optogenetically evoked, asynchronous Sr^2+^-oEPSC in iSPNs fell during the on-state and rose during the off-state, consistent with the well described role of D2Rs in the modulation of long-term synaptic plasticity in these cells ^56,63,86,87^. Importantly, the frequency of asynchronous Sr^2+^-oEPSCs did not change between these states, arguing that the total number of synapses was unchanged. Although the augmentation of synaptic strength in the off-state was consistent with previous studies ^41^, the apparent stability in number of synapses was not.

Previously, it was reported that iSPN spine density – and by inference synaptic density – increased in the off-state after the induction of dyskinesia ^40,42,66^. Although our estimates of spine density using 2PLSM rose in the off-state, reaching values close to those seen prior to 6-OHDA lesioning, higher-resolution confocal imaging of iSPN dendrites found only a modest change in spine density, well below values seen in controls. Taken together, our studies suggest that there is an alternative interpretation of the data. With the best optical approaches, the density of SPN spines in proximal dendrites is roughly 2.6 spines/µm, with many spines being less than a half micron in diameter ^67,68^. Using confocal microscopy with a high numerical aperture (NA) lens (1.49), proximal spine density was estimated to be about 2 spines/µm, suggesting that about a quarter of the spines were being missed. Using 2PLSM with a lower NA lens (0.9), proximal spine density was estimated to be about 1.3 spines/µm, suggesting that roughly half of the spines were not being detected. The most parsimonious interpretation of the imaging and physiology is that what is changing between on- and off-states is not the number of spines, but rather their diameter and resolvability with optical methods. Indeed, when the confocal analysis was limited to large spines (>0.4 µm diameter) (Supplemental Fig. S5), there was good alignment with the 2PLSM data. This is an important point, because it suggests that iSPNs in an adult brain are not breaking and forming new synapses, but rather, are modulating their strength (which results in a size change). It is still the case, of course, that this modulation in synaptic strength is uncoupled from actions and their outcomes, unlike the situation in the normal striatum. As noted above, this should result in a degradation in synaptic information and an impaired ability to properly coordinated movement.

### Dysregulation of ChIs contributed to LID severity

In the healthy striatum, the interaction between DA and ACh release by ChIs is important in modulating the activity of SPNs and motor control ^15,88^. In both dSPNs and iSPNs, this interaction is largely antagonistic, which creates a means of dynamically regulating basal ganglia output. In iSPNs, DA activation of D2Rs diminishes somatodendritic excitability and promotes LTD at glutamatergic synapses, whereas ACh activation of M1Rs does the opposite; in dSPNs, D1Rs promote somatodendritic excitability and LTP at glutamatergic synapses whereas M4Rs do the opposite ^20,23,24,48,56,84,85^. Complementing this interaction at the level of SPNs, ACh triggers DA release by activating nicotinic receptors on dopaminergic terminals ^89–91^, whereas dopaminergic activation of D2Rs expressed by ChIs inhibits the autonomous spiking of ChIs and ACh release from axon terminals ^15–18,27^.

In PD, this dynamic interaction is disrupted. The loss of the striatal dopaminergic innervation in PD patients has long been thought to dis-inhibit ChIs, leading to a hyper-cholinergic state, a persistent imbalance in the excitability of iSPNs and dSPNs and ultimately bradykinesia ^1,21,27–33^. Our experiments using the genetically encoded optical sensor of ACh (GRAB_ACh3.0_) are consistent with this hypothesis, showing that in the 6-OHDA lesioned striatum, electrically evoked ACh release is significantly greater than in control striata. This up-regulation was primarily attributable to the loss of presynaptic D2R signaling, as the D2R agonist quinpirole was able to almost eliminate evoked ACh release. That said, the fact that peak ACh release was significantly greater in the lesioned striatum than in the control striatum in the presence of the D2R antagonist sulpiride, also argues that mAChR autoreceptor function was impaired by DA depletion, as previously reported ^27^.

Although the role of elevated ACh release by ChIs in parkinsonism is widely accepted and consistent with the clinical utility of mAChR antagonists, the involvement of ChIs in LID has been controversial ^34–36^. The issue in the field is not whether they influence LID severity, but rather how. Several lines of study suggest that hyper-activity of ChIs exacerbates LID severity. For example, ablation or chemogenetic inhibition of ChIs lessens the severity of LID in rodent models ^38,39^, as does genetic perturbation of ChIs ^92^. On the other hand, enhancing M4R signaling in dSPNs attenuates LID induction ^20,37^ and genetic deletion of D5 DA receptors from ChIs, which appears to lower their excitability, worsens LID ^93^. Similarly, as shown here, deleting M1Rs in iSPNs attenuated LID induction, as did deletion of its downstream signaling partner, CDGI.

Much of this controversy may stem from a failure to distinguish between what is happening in on- and off-states. Our GRAB_ACh3.0_ experiments clearly show that DA continues to robustly inhibit ACh release from ChIs in the striatum of dyskinetic mice. As a consequence, it seems very unlikely that when striatal DA levels rise during the on-state that ChI ACh release rises in parallel, regardless of whether there are modest increments in ChI intrinsic excitability induced by repeated levodopa treatment ^79,94^. In contrast, during the off-state, when striatal DA levels plummet, ChIs are undoubtedly disinhibited. Off-state ACh release by ChIs also should be augmented by an elevation in intrinsic excitability induced by repeated levodopa treatment ^79,94^. The elevation in the excitability of iSPNs in the off-state is precisely what would be predicted by an up-regulation in ACh release and activation of M1Rs ^22–24,85^. The off-state reversal of the on-state elevation in dSPN excitability is also consistent with engagement of M4Rs ^20,21,37^. The analysis of SPN synaptic plasticity also aligned with this hypothesis; that is, during the off-state, the strength of iSPN glutamatergic synapses rose, whereas synaptic strength fell in dSPNs. These synaptic shifts are consistent with the well-described effects of ACh (and DA) on synaptic plasticity in these two cell types ^9,10,34,70,71^. This functional plasticity was mirrored structurally, as the number of spines that were discernible with 2PLSM rose in iSPNs and fell in dSPNs during the off-state.

Taken together, these data are consistent with the conclusion that ChIs play an important role in the pathophysiology underlying LID – in agreement with previous work. Further evidence for this conclusion comes from the demonstration that disrupting M1R signaling in iSPNs, either by deleting CDGI globally or by CRISPR-Cas9-mediated deletion of *Chrm1* specifically from them, blunted the induction of LID. However, our results suggest that the engagement of ChIs and their remodeling of striatal circuitry is occurring primarily during the off-state. The importance of off-state striatal adaptations is underscored by recent work showing that during this period, iSPNs up-regulate the expression of GluN2B-containing NMDA receptors, which enable the induction of LTP ^62^. This observation is consistent with the off-state strengthening of corticostriatal synapses and spine enlargement in iSPNs reported here. Furthermore, the GluN2B up-regulation is necessary for not just the induction of LID, but its expression after repeated levodopa treatment ^62^.

What is less clear is how the off-state adaptations in iSPNs promote dyskinesia with the next dose of levodopa. Previous work has shown that enhancing the intrinsic excitability of iSPNs in the on-state attenuates LID ^61^. Hence, had the elevation in off-state iSPN excitability persisted into the on-state, it would have been anti-dyskinetic. But this alteration was not persistent and reversed after levodopa treatment. As discussed above, the mechanisms responsible for the reversal remain to be determined. Nevertheless, increased excitability in the off-state, particularly in dendrites, will enhance the induction of postsynaptic LTP in iSPNs, which depends upon depolarization and NMDAR opening. Indeed, deletion of LTP-promoting ^22^ M1Rs (or their dendritic signaling partner CDGI), blunted the off-state strengthening of corticostriatal synapses. How might the induction of synaptic potentiation in iSPNs during the off-state promote LID? Appropriately timed and coordinated movement requires co-activation of iSPN and dSPN ensembles. The architecture of SPN ensembles is largely dependent upon how the activity in cortical ensembles is translated by striatal circuits. There are a variety of factors that contribute to this translation, but the weighting of synaptic inputs by SPNs is certainly an important one. This synaptic weight or strength is normally shaped by experience and repetition ^95–97^. This is the basis for goal-directed and habit learning based in striatal circuits. However, in the parkinsonian striatum, this linkage is lost. Recent work has underscored the importance of aberrant learning in PD, suggesting that engagement of cortical circuitry associated with a particular action in the absence of striatal DA (and elevation in ACh) contributes to the inability to perform that action in the future ^98^. It is easy to imagine that in this situation, the striatal circuitry ‘interprets’ the neuromodulator imbalance as a negative outcome, leading to strengthening of connectivity between action-associated cortical circuits and iSPN ensembles, while weakening those with dSPN ensembles.

This kind of aberrant, outcome independent striatal learning could also be happening in the off-state. If this were the case, the identity of the cortical synapses being strengthened would depend upon which cortical ensembles were active in the off-state, as coincident pre- and post-synaptic activity is necessary for LTP induction ^56^. The identity of these ensembles is uncertain, especially given the profoundly hypokinetic nature of the off-state. Conversely, the sustained elevation in striatal DA and drop in ACh during the on-state also creates an aberrant learning signal in the striatum, akin to a positive outcome for any and all actions, promoting corticostriatal LTD in iSPNs ^56,63,86,87^. This form of plasticity also requires coordination of pre- and post-synaptic signaling that engages group I metabotropic glutamate receptors (mGluRs) ^86^; mGluR signaling in other neurons is enhanced by GluN2B-containing NMDARs ^99,100^, raising the possibility that their augmentation during the off-state sets the stage for more robust, aberrant LTD during the on-state. This shift could be similar to a form of metaplasticity seen in a variety of other neurons after bouts of sustained activity ^101–103^. Regardless, this form of aberrant ‘learning’ could make it very difficult to activate iSPN ensembles that suppress unwanted movement, contributing to the pathway imbalance underlying dyskinetic behavior.

### Translational implications

At present, the strategies for alleviating LID in PD patients are very limited. Our studies demonstrate that suppressing M1R signaling in iSPNs alleviates LID in a rodent model. Global antagonism of M1Rs would come with unacceptable side effects. The best translational path forward for this observation would involve a regionally targeted gene therapy. The recent development of enhancer sequences that can be used to limit gene expression specifically to iSPNs in the dorsal striatum ^104^, creates a potential path forward for a novel gene therapy targeting M1Rs or CDGI that does not rely upon neurosurgical approaches.

## Acknowledgements

We thank the members of the Surmeier lab for their support and comments on the manuscript. Special thanks go to Sasha Ulrich and Milos Aleksic for their expert technical support with the behavioral experiments. This work is supported by the JPB foundation (to D.J.S.), NS34696 (to D.J.S.), William N. and Bernice E. Bumpus Foundation (to D.J.S, S.Z. and A.M.G.), Aligning Science Across Parkinson’s [ASAP020551] through the Michael J. Fox Foundation for Parkinson’s Research (MJFF) (to D.J.S.), Jim & Joan Schattinger (to A.M.G.) and Mr. Robert Buxton (to A.M.G.).

## Author Contributions

S.Z. and D.J.S. designed the experiments. S.Z. performed all electrophysiological, two-photon imaging, and immunohistochemistry experiments and data analysis (with D.W.’s help). Q.C. performed sparse labeling-enabled confocal reconstruction of dendritic structure. S.Z. and L.S. performed the behavioral experiments and analysis. J.R.C. and A.M.G. generated the CDGI KO mice and discussed strategies. T.T. generated the M1R CRISPR constructs. S.Z. and D.J.S. prepared the manuscript with edits by A.M.G.. All authors approved the final version.

## Availability Statement

The data, code, protocols, and key lab materials used and generated in this study are listed in a Key Resource Table alongside their persistent identifiers at www.doi.org/10.5281/zenodo.14589126.

## Detailed Methods

### Animals

Animal use procedures were approved by the Northwestern Institutional Animal Care and Use committee. Male C57Bl/6 hemizygous mice (7-12 weeks of age) expressing tdTomato or eGFP under control of Drd1a or Drd2 receptor regulator elements (RRID: MMRRC_030512-UNC and RRID: MMRRC_000230-UNC, backcrossed to C57BL/6 background) were used. In some experiments, these mice were crossed with CDGI KO mice in a C57BL/6 J background originally generated at MIT ^22,25^. BAC transgenic mice expressing Cre recombinase under control of the A2aR regulatory elements (RRID:MMRRC 031168-UCD) were used for high-resolution confocal microscopy and M1R CRISPR experiments.

### Unilateral 6-OHDA model of PD and LID

Mice were anesthetized using an isoflurane precision vaporizer (at 5% isofluorane during induction and 2% isofluorane during maintenance phase) and positioned in a stereotaxic frame (David Kopf Instruments, model 940). Mice were administered with analgesics meloxicam (METACAM®, 0.1mg/kg, s.c., Covetrus) before surgery. After the skin and fascia were retracted to reveal the skull, a small hole was drilled over the MFB. 3.5 mg ml^−1^ free base 6-OHDA hydrochloride (Sigma Aldrich, Cat. No. H4381) freshly dissolved in saline with 0.02% L-ascorbic acid (Sigma Aldrich, Cat. No. A92902) was injected using a glass calibrated micropipette (Drummond Scientific Company, Broomall, PA, 200005) at the following coordinates: AP (mm relative to Bregma): –0.7; ML: –1.2; DV: –4.75 (from dura). Mice were monitored daily post-op and supplemented with saline injections and high fat/high sucrose food as needed ^62^. Three to four weeks after surgery, the degree of lesioning of nigrostriatal DA neurons was assessed with a drug-free cylinder test ^40,62^. Within 1-7 days of the cylinder test, mice underwent behavioral testing for abnormal involuntary movements (AIMs) as previously described ^4,40,62^. In brief, mice were transferred to a behavioral testing room, placed in clean cage bottoms without bedding, and administered intraperitoneal injections of L-DOPA at 6 mg/kg for the first two sessions and 12 mg/kg for the later sessions. Benserazide was co-administered at 12 mg/kg to inhibit peripheral conversion of levodopa to DA. Behavioral testing was performed every other day for a total of five test sessions. AIMs (axial, limb, and orolingual movements) were scored as previously described ^4,40,62^: abnormal axial, limb, and orolingual behaviors were observed for one minute every 20 min and rated on a scale from 0-4 for each parameter on the basis of duration and continuity. Mice were sacrificed for *ex vivo* experiments 24-48 hours (off-state) or 0.5 hour (on-state) after the last levodopa administration. Striatal and nigral sections from a subset of mice were stained with tyrosine hydroxylase to verify successful lesion.

### Immunohistochemistry and confocal microscopy

Anesthetized mice were perfused transcardially with saline briefly (∼1 min) and then with ice-cold 4% paraformaldehyde (wt/vol) in 1x phosphate buffered saline (4% PFA-PBS). Brains were dissected out and post-fixed in 4% PFA-PBS overnight at 4°C. Coronal slices (100 μm-thick) were cut using a vibratome (Leica VT1200), permeabilized and blocked at 4°C in PBS including 5% normal goat serum and 0.2% Triton-X100 (NGS/PBST), incubated with rabbit phospho-ERK antibody (Cell Signaling, #9101; 1:100 dilution) and mouse tyrosine hydroxylase antibody (Immunostar, #22941; 1:200 dilution) diluted in NGS-PBST overnight at 4°C. After four washes in NGS/PBST, the slices were incubated with 1:1000 goat anti-rabbit Alexa Fluor 488 or goat anti-mouse Alexa Fluor 555 (Invitrogen A-11001 and A-21428) for 2 hours at room temperature. Slices were washed with NGS-PBST four times and PBS once, mounted with VECTASHIELD Antifade Mounting Medium (Vector Laboratories, H-1000-10), and viewed under an automated laser scanning confocal microscope (FV10i-DUC; Olympus). Images were adjusted for brightness, contrast, and pseudo-coloring in ImageJ (US National Institutes of Health).

### Viral injections

To study synaptic responses at corticostriatal synapses, 0.15 μl AAV5-hSyn-hChR2(H134R)-eYFP (Addgene #26973) was injected into the M1 motor cortex ipsilateral to the lesion site at the following coordinate (mm relative to Bregma): ML: -1.60, AP: 1.15, DV: 1.55. To sparsely label iSPNs for high-resolution confocal imaging of dendritic spines, *Adora2*-Cre mice (∼2 months old) were injected with AAV9-pCAG-flex-eGFP-WPRE (Addgene #51502) (0.450 μl, titer of 5 x 10^10^ viral genome/ml) into the left DLS at (mm relative to Bregma): AP: 0.7, ML: -2.35, DV: ∼3.35. To express the ACh sensor GRAB_ACh3.0_ in the striatum, 0.6 μl of AAV5-hSyn-ACh3.0(ACh4.3) (WZ Biosciences, YL001003-AV5) was injected into the DLS ipsilateral to the 6-OHDA lesion at the following coordinate (mm relative to Bregma): ML: -2.2, AP: 0.8, DV: 3.3. For M1R gene deletion using CRISPR-Cas9, well-lesioned animals were randomly assigned to receive viral injection of either a mixture of two viruses (at 1:1 ratio): AAV9-hSyn-DIO-saCAS-minWPRE and AAV9-U6-gRNA(M1R)-U6-gRNA2-hSyn-FusionRed (*M1R CRISPR*, custom made by Virotek), or a mixture of saline and the gRNA virus (*control*). The viral injection (0.5 μl) was in the DLS (ipsilateral to the 6-OHDA lesion) at a total of 6 sites at the following coordinates (mm relative to Bregma): AP: 0.9, ML: –2.3, DV: –3.4 and −2.8; AP: 0.6, ML: –1.5, DV: –3.4 and −2.8; AP: 0.24, ML: –1.9, DV: –3.3 and −2.7. For *ex vivo* experiments that required identification of CRISPR-expressing iSPNs, a third virus: AAV9-EF1a-DIO-eYFP (Addgene #27056) was mixed with M1R CRSIPR or control (at 1:1:1 ratio) and injected. All experiments were performed about 3 weeks (for Cre-independent expression) or 4-5 weeks (for Cre-dependent expression) after viral injections.

### Slice electrophysiology

Mice were deeply anesthetized with a mixture of ketamine (100 mg/kg) and xylazine (7 mg/kg) and perfused transcardially with ice-cold sucrose-based cutting solution containing (in mM): 181 sucrose, 25 NaHCO_3_, 1.25 NaH_2_PO_4_, 2.5 KCl, 0.5 CaCl_2_, 7 MgCl_2_, 11.6 sodium ascorbate, 3.1 sodium pyruvate and 5 glucose (305 mOsm/l). Sagittal slices (280-μm thick) were sectioned using a vibratome (Leica VT1200). After cutting, slices were incubated at 34 °C for 30 min in ACSF containing (in mM): 124 NaCl, 3 KCl, 1 NaH_2_PO_4_, 2.0 CaCl_2_, 1.0 MgCl_2_, 26 NaHCO_3_ and 13.89 glucose, after which they were stored at room temperature until recording. External solutions were oxygenated with carbogen (95%CO_2_/5%O_2_) at all time.

Individual slices were transferred to a recording chamber and continuously superfused with ACSF (2-3 ml/min, 31-32°C). D1-Tdtomato- or D2-eGFP-expressing SPNs in the striatum were first identified with an Olympus BX-51-based two-photon laser scanning microscope (Ultima, Bruker). Whole-cell patch clamp was then performed in identified SPNs, aided by visualization with a 60X/0.9NA water-dipping objective lens and a ½” CCD video camera (Hitachi) imaged through a Dodt contrast tube and a 2x magnification changer (Bruker). For somatic, dendritic and morphological experiments, patch pipettes (3-4 MΩ resistance) were loaded with internal solution containing (mM): 115 K-gluconate, 20 KCl, 1.5 MgCl_2_, 5 HEPES, 0.2 EGTA, 2 Mg-ATP, 0.5 Na-GTP, 10 Na-phosphocreatine (pH 7.25, osmolarity 280-290 mOsm/L). Cells were recorded in the current-clamp configuration. For Sr^2+^-oEPSC experiments, patch pipettes (3-4 MΩ resistance) were loaded with 120 CsMeSO_3_, 5 NaCl, 0.25 EGTA, 10 HEPES, 4 Mg-ATP, 0.3 Na-GTP, 10 TEA, 5 QX-314 (pH 7.25, osmolarity 280-290 mOsm/L). SPNs were held at -70 mV in the voltage-clamp configuration. After patching, recording solution was changed to Ca^2+^-free ACSF containing 3 mM SrCl_2_ and 10 μM gabazine (10 μM, to suppress GABA_A_-mediated currents). Slices were incubated with this Ca^2+^-free solution for 25 min before recording. EPSCs were evoked every 30 s by whole-field LED illumination (single 0.3-ms pulses). All the electrophysiological recordings were made using a MultiClamp 700B amplifier (Axon Instrument, USA), and signals were filtered at 2 kHz and digitized at 10 kHz. Voltage protocols and data acquisition were performed by PraireView 5.3 (Bruker). The amplifier command voltage and all light source shutter and modulator signals were sent via the PCI-NI6713 analog-to-digital converter card (National Instruments, Austin, TX).

### Imaging of spines and dendritic Ca^2+^ transients with two-photon laser scanning microscopy (2PLSM)

The pipette solution (EGTA omitted) was supplemented with 100 μM Fluo-4 (Thermo Fisher Scientific, F14200) and 50 μM Alexa Fluor 568 hydrazide (Thermo Fisher Scientific, A10437). After whole-cell recording configuration was established, cells were allowed to equilibrate with dyes for at least 15 min before imaging. The recorded SPN was visualized using 810 nm excitation laser (Chameleon Ultra II, Coherent, Santa Clara, USA). Dendritic structure was visualized by the red signal of Alexa Fluor 568 detected by a Hamamatsu R3982 side-on photomultiplier tube (PMT, 580-620 nm). Calcium transients, as signals in the green channel, were detected by a Hamamatsu H7422P-40 GaAsP PMT (490-560 nm, Hamamatsu Photonics, Japan). Signals from both channels were background subtracted before analysis. Line scan signals were acquired with 128 pixels per line resolution and 10 μs/pixel dwell time along a dendritic segment. Ca^2+^ signals were quantified as the area of increase in green fluorescence from baseline normalized by the average red fluorescence (ΔG/R). Dendritic excitability index was measured as the ratio of the Ca^2+^ signal from a distal location to the Ca^2+^ signal from a proximal location on the same dendrite. Only data with similar baseline levels (G_0_/R_0_) for proximal and distal locations were included.

For assessment of dendritic spine density, images of dendritic segments (proximal: ∼40 μm from soma; distal: > 80 μm from soma) were acquired with 0.15 μm pixels with 0.3 μm z-steps. Images were deconvolved in AutoQuant X3.0.4 (MediaCybernetics, Rockville, MD) and semi-automated spine counting was performed using 3D reconstructions in NeuronStudio (CNIC, Mount Sinai School of Medicine, New York). On average, two proximal and two distal dendrites were imaged and analyzed per neuron.

### Two-photon imaging of ACh sensor

ACh release was assessed by imaging GRAB_ACh3.0_, a genetically encoded fluorescent sensor of ACh ^72^, using 2PLSM. Acute slices with striatal expression of GRAB_ACh3.0_ were prepared as described above, transferred to a recording chamber, and continuously perfused with normal ACSF at 32– 34°C. A two-photon laser (Chameleon Ultra II, Coherent, Santa Clara, CA) tuned to 920 nm was used to excite GRABACh3.0. Fluorescence was imaged using an Ultima In Vitro Multiphoton Microscope system (Bruker, Billerica, MA) with an Olympus 60x/0.9 NA water-immersion objective lens and a Hamamatsu H7422P-40 GaAsP PMT (490 nm to 560 nm, Hamamatsu Photonics, Hamamatsu, Japan). Time series images of the GRAB_ACh3.0_ were acquired with 0.388 µm × 0.388 µm pixels, 8-µs dwell time and a frame rate of 21.26 fps. After 3-s baseline acquisition, synchronous ACh release was evoked by delivering a single (1 ms x 0.3 mA) or a train of 20 electrical stimuli (1 ms x 0.3 mA at 20 Hz) by a concentric bipolar electrode (CBAPD75, FHC) placed at 200 μm ventral to the region of interest. Imaging was continued for at least another 5 s. Two trials were performed for each stimulation protocol and data averaged. The slices were imaged for (in this sequence) control, 50 nM DA (only for lesioned mice), 10 μM quinpirole and 10 μM sulpiride treatment conditions, with at least 5 min of perfusion for each treatment. The minimal (F_min_) and maximal fluorescence intensity (F_max_) were determined by applying 10 µM TTX (to block any basal transmission) and 100 µM acetylcholine chloride (to saturate GRAB_ACh3._0 signal), respectively. The whole image was the region of interest (ROI) used for analysis. Fluorescent intensity data were analyzed by custom Python code (accessible upon request). Briefly, the fluorescence intensity values were first background-subtracted (the background resulted from PMT was measured by imaging with same PMT voltage but zero laser power). Baseline fluorescence F_0_ was the average fluorescence over the 1 s period right before stimulation. ΔF = F - F_0_ was normalized by (F_max_ − F_min_) and then analyzed.

### High-resolution confocal microscopy of sparsely labeled neurons

*Adora2*-Cre mice injected with Cre-dependent eGFP virus and treated with 4 different conditions (control, 6-OHDA, LID off-state, LID on-state) were anesthetized with isoflurane followed by ketamine/xylazine mixture. LID on-state mice received a sixth dose of levodopa (12 mg/kg, supplemented with 12 mg/kg benserazide) one hour before anesthesia. The mice were transcardially perfused with 1x phosphate buffered saline (PBS, ∼20 ml) followed by 4% paraformaldehyde (PFA, ∼30 ml). Brains were then dissected out, postfixed in 4% PFA for 1.5-2hrs, transferred to 1x PBS with 0.1% sodium azide and stored at 4°C until sectioning. 60-μm sagittal sections containing the DLS were cut using a Leica VT1200S vibratome. Sections were mounted onto glass slides with No. 1.5 cover glasses using ProLong™ Diamond Antifade Mountant (Thermo Fisher Scientific, Cat. No. P36961).

A Nikon AXR confocal laser microscope from Center for Advanced Microscopy & Nikon Imaging Center at Northwestern University was used to image dendritic spines from sparsely labeled iSPNs in fixed brain sections. The z-stack images were acquired with a 60x oil immersion objective (NA = 1.49) at 0.125 μm intervals with 0.09 μm pixel size. The images were then de-noised and deconvolved with Nikon NIS-Elements AR 5.41.02 software. About 30 μm dendritic segments measuring at ∼30 μm from the soma were analyzed. Spine density analysis was performed with Imaris 10.0.0 software (Oxford Instruments). Briefly, the dendrite of interest was first isolated by adding a new Surface. ‘Labkit for pixel classification’ was adopted to train the detection of signals over noises to create a tight surface. The surface was further cut to only include the dendritic segment of interest. Filament for the dendritic segment was then created with the aid of an embedded supervised learning function. Spine detection was subsequently performed with the thinnest spine head set at 0.188 μm, maximum spine length set at 5 μm and no branched spines were allowed. Spines were further determined with an embedded supervised learning function and manually corrected if necessary.

### Data acquisition, analysis and statistics

Electrophysiology and imaging data were acquired using PCI-NI6052E analog-to-digital converter card (National Instruments) and *Praire View* 5.3 (Bruker). Off-line analyses of electrophysiology data (except Sr^2^-oEPSCs), calcium imaging data and time-series imaging data were performed using custom-written Python scripts (available upon request). Amplitudes of Sr^2^-oEPSCs were analyzed automatically using TaroTools Event Detection in Igor Pro 8 (WaveMetrics, Portland, Oregon) followed by visual verification. Events were measured between 40 ms to 400 ms after photostimulation. The threshold for detection of an event was greater than 5 SD above the noise. All summary data were presented as non-parametric box-whisker plots and statistical analyses performed by Prism 6 (GraphPad). The stated *n* indicates the number of cells (in electrophysiology experiments) or the number of mice (in AIMs testing). Comparisons were made using Mann-Whitney test (unpaired) or Wilcoxon test (paired) and differences with p < 0.05 are considered statistically significant.

---

**Supplemental Figure 1.**
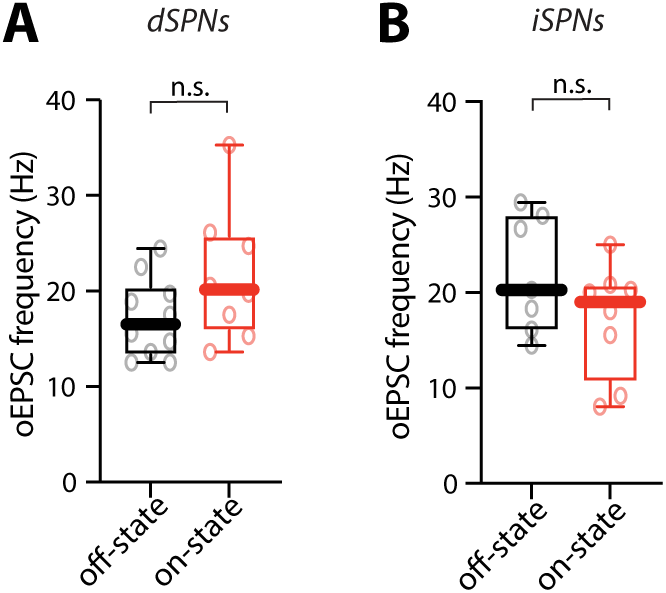
The frequency of Sr^2+^-oEPSC was unaltered between off- and on-states in both dSPNs and iSPNs. (A) Box plot summary of Sr^2+^-oEPSC frequency in off- and on-state dSPNs (off-state, n = 10 cells from 4 mice; on-state, n = 8 cells from 4 mice). n.s., not statistically significant, Mann-Whitney test. (B) Box plot summary of Sr^2+^-oEPSC frequency in off-state and on-state iSPNs (off-state, n = 7 cells from 4 mice; on-state, n = 8 cells from 4 mice). n.s., not statistically significant, Mann-Whitney test.

**Supplemental Figure 2.**
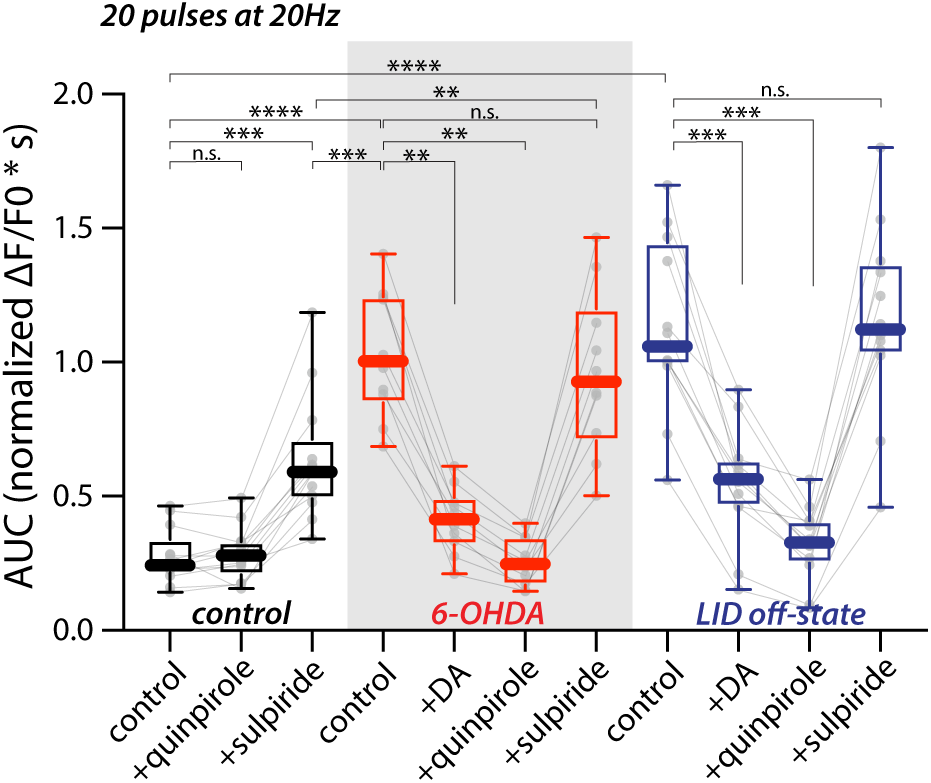
ACh release evoked by burst stimulation was elevated in 6-OHDA lesioned and LID off-state mice and suppressed by D2R agonists Box plot summary of GRAB_ACh3.0_ signals evoked by burst stimulation in unlesioned, 6-OHDA lesioned and LID off-state mice. Same as with single stimulation, ACh release evoked by burst stimulation (20 pulses at 20 Hz) was significantly elevated in 6-OHDA lesioned and LID off-state mice. Bath application of DA (50 nM) or quinpirole (10 μM) strongly suppressed ACh release (unlesioned, n = 13 ROIs from 3 mice; 6-OHDA, n = 10 ROIs from 3 mice; off-state, n = 12 ROIs from 5 mice). **** p < 0.0001, *** p < 0.001, ** p < 0.01, n.s., not statistically significant, Mann-Whitney test for unpaired data and Wilcoxon for paired data.

**Supplemental Figure 3.**
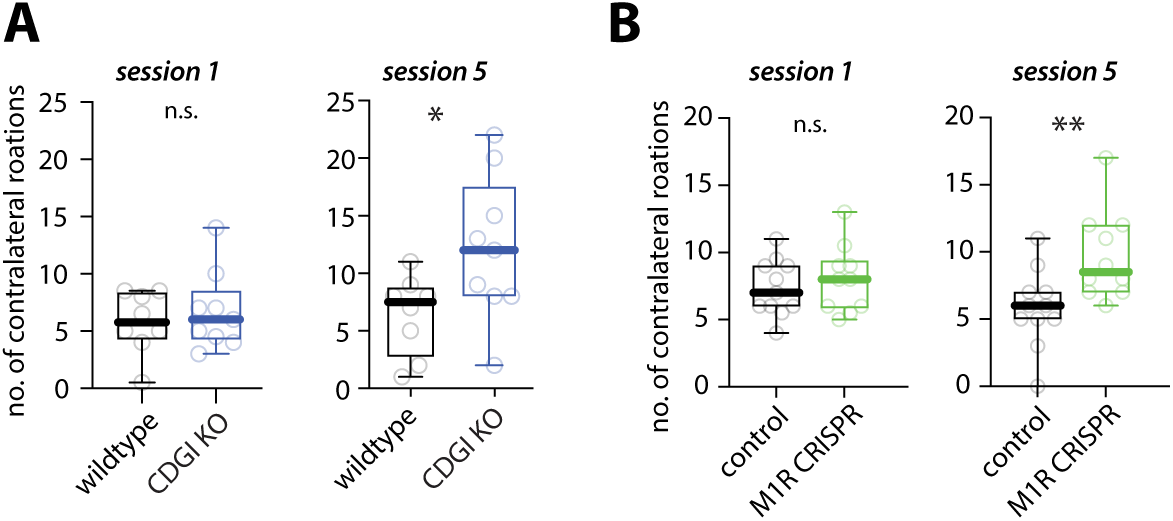
Genetic perturbation of M1R-CDGI signaling in iSPNs increases the therapeutic effect of levodopa in mice with established LID. (A) Box plot summary of the number of contralateral rotations (in 30 s) recorded 40 min after the first and fifth levodopa administration in wildtype or CDGI KO mice (wildtype n = 8 animals; CDGI KO, n = 9 animals). * p < 0.05, n.s. not statistically significant, Mann-Whitney test. The session 5 data were the same as in Fig. 7I. (B) Box plot summary of the number of contralateral rotations (in 30 s) recorded 40 min after the first or fifth levodopa administration in control or M1R CRISPR mice (control n = 11 animals; M1R CRISPR, n = 10 animals). ** p < 0.01, n.s. not statistically significant, Mann-Whitney test.

**Supplemental Figure 4.**
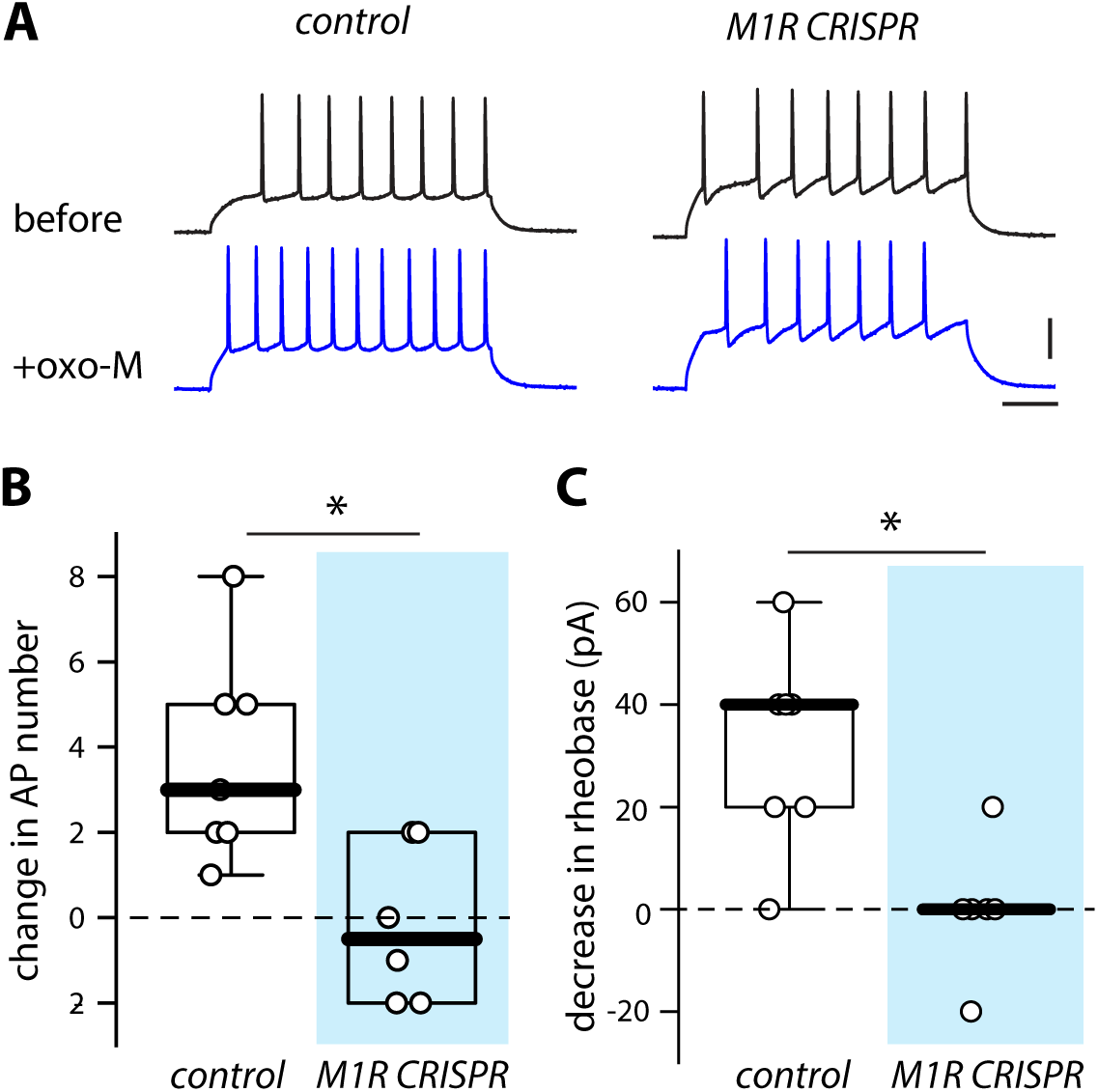
Functional validation of M1R CRISPR. (A) Example traces of somatic voltage recordings in response to a 180-pA current injection from iSPNs expressing M1R CRISPR or gRNA alone (control) before and after application of oxo-M. Scale bars are 40 mV and 100 ms. (B) Box plot summary of the effect of oxo-M on the number of action potentials (APs) evoked (control, n = 7 cells from 3 mice; M1R CRISPR, n = 6 cells from 3 mice). The increase in somatic excitability by oxo-M was prevented by M1R CRISPR. * p < 0.05, Mann-Whitney test. (C) Box plot summary of the effect of oxo-M on rheobase. * p < 0.05, Mann-Whitney test.

**Supplemental Figure 5.**
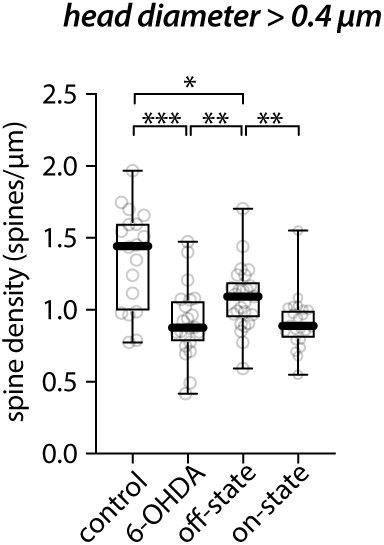
Density of spines with larger head diameters detected by high-resolution confocal microscopy Box plot summary of the density of spines with > 0.4 μm head diameter in proximal dendrites of sparsely labeled iSPNs imaged by high-resolution confocal microscopy (control: n = 18 dendrites from 4 mice; 6-OHDA: n = 22 dendrites from 3 mice; off-state: n = 25 dendrites from 4 mice; on-state: n = 22 dendrites from 4 mice). This result was similar to the total spine density detected by 2PLSM (Fig. 4B). *** p < 0.001, ** p < 0.01, * p < 0.05, Mann-Whitney test.

